# Crosstalk between regulatory elements in the disordered TRPV4 N-terminus modulates lipid-dependent channel activity

**DOI:** 10.1101/2022.12.21.521430

**Authors:** Benedikt Goretzki, Christoph Wiedemann, Brett A. McCray, Stefan L. Schäfer, Jasmin Jansen, Frederike Tebbe, Sarah-Ana Mitrovic, Julia Nöth, Jack K. Donohue, Cy M. Jeffries, Wieland Steinchen, Florian Stengel, Charlotte J. Sumner, Gerhard Hummer, Ute A. Hellmich

**Affiliations:** Friedrich Schiller University Jena, Faculty of Chemistry and Earth Sciences, Institute of Organic Chemistry and Macromolecular Chemistry, Humboldtstraße 10, 07743 Jena, Germany; Centre for Biomolecular Magnetic Resonance (BMRZ), Goethe University, Max-von-Laue Str. 9, 60438 Frankfurt am Main, Germany; Department of Neurology, Johns Hopkins University School of Medicine, Baltimore, MD, USA; Department of Theoretical Biophysics, Max Planck Institute of Biophysics, Max-von-Laue Str. 3, 60438 Frankfurt am Main, Germany; Department of Biology, University of Konstanz, Universitätsstraße 10, 78457 Konstanz, Germany; Konstanz Research School Chemical Biology, University of Konstanz, Universitätsstraße 10, 78457 Konstanz, Germany; Department of Chemistry, Section Biochemistry, Johannes Gutenberg-University Mainz, Joachim-Becher-Weg 30, 55128 Mainz, Germany; European Molecular Biology Laboratory, EMBL Hamburg Unit, Deutsches Elektronen-Synchrotron, Notkestraße 85, 22607 Hamburg, Germany; Center for Synthetic Microbiology (SYNMIKRO) & Department of Chemistry, Philipps-University Marburg, Karl-von-Frisch-Str. 14, 35043 Marburg, Germany; Department of Neuroscience, Johns Hopkins University School of Medicine, Baltimore, MD, USA; Institute of Biophysics, Goethe University Frankfurt, Max-von-Laue Str. 1, 60438 Frankfurt am Main, Germany

**Keywords:** TRP channel, intrinsically disordered region, long-range intramolecular communication, PIP_2_ binding, integrated structural biology, MD simulations

## Abstract

Intrinsically disordered regions (IDRs) are essential for membrane receptor regulation but often remain unresolved in structural studies. TRPV4, a member of the TRP vanilloid channel family involved in thermo- and osmosensation, has a large N-terminal IDR of approximately 150 amino acids. With an integrated structural biology approach, we analyze the structural ensemble of the TRPV4 IDR and identify a network of regulatory elements that modulate channel activity in a hierarchical lipid-dependent manner through transient long-range interactions. A highly conserved autoinhibitory patch acts as a master regulator by competing with PIP_2_ binding to attenuate channel activity. Molecular dynamics simulations show that loss of the interaction between the PIP_2_-binding site and the membrane reduces the force exerted by the IDR on the structured core of TRPV4. This work demonstrates that IDR structural dynamics are coupled to TRPV4 activity and highlights the importance of IDRs for TRP channel function and regulation.

## Introduction

The majority of eukaryotic ion channels contain intrinsically disordered regions (IDRs), which play important roles in protein localization, channel function and the recruitment of regulatory interaction partners^1–3^. In some transient receptor potential (TRP) channels, IDRs make up more than half of the entire protein sequence^4^. Among the mammalian TRP vanilloid (TRPV) subfamily, TRPV4 has the largest N-terminal IDR, ranging from ~130 to ~150 amino acids in length depending on the species^4–6^. TRPV4 is a Ca^2+^-permeable plasma membrane channel that is widely expressed in human tissues. It is remarkably promiscuous, and stimuli include pH, moderate heat, osmotic and mechanic stress, and various chemical compounds^7,8^. TRPV4 also garnered attention due to the large number of disease-causing mutations with distinct tissue-specific phenotypes primarily affecting the nervous and skeletal systems^9–13^. Among others, roles in cancer as well as viral and bacterial infections have also been described^14–16^.

Crystal structures of the isolated TRPV4 ankyrin repeat domain (ARD) were among the first regions of a TRP channel to be resolved, showing a compact, globular protein domain with six ankyrin repeats^10,17,18^. Together with the IDR, the ARD forms the channel’s cytoplasmic N-terminal domain (NTD). Furthermore, near full-length frog and human TRPV4 cryo-EM and X-ray crystallography structures are available, but lack the IDR, which was partially or fully deleted to facilitate structure determination^19,20^. Short stretches of N- and C-terminal IDRs were found previously to interact with the ARD in TRPV2 and TRPV3 cryo-EM structures^21–23^, but no complete TRP(V) channel IDR has been visualized to date.

The TRPV4 NTD is responsible for channel sensitivity to changes in cell volume^24^, its reaction to osmotic and mechanical stimuli^25,26^ and the interaction with regulatory binding partners^27–30^. Therefore, a structural characterization of the TRPV4 NTD including its large IDR is critical to understanding TRPV4 regulation in detail.

To date, two regulatory elements in the N-terminal TRPV4 IDR have been described: (i) a proline-rich region directly preceding the ARD that enables protein-dependent channel desensitization^27,28,30^; and (ii) a phosphatidylinositol-4,5-bisphosphate (PIP_2_)-binding site composed of a stretch of basic and aromatic residues directly N-terminal to the proline-rich region^31^. PIP_2_ is a plasma membrane lipid and an important ion channel regulator^32,33^. In TRPV4, mutation of the PIP_2_-binding site abrogates PIP_2_-dependent channel sensitization in response to osmotic and thermal stimuli^31^. It remains unknown whether the TRPV4 IDR contains additional regulatory elements and how they may mediate channel regulation. An understanding of the dynamic properties of a complete TRP channel IDR at atomic resolution is currently lacking, which complicates the search for such putative regulatory elements and their structural crosstalk.

Here, we used an integrated structural biology approach to analyze the structural ensemble of the TRPV4 N-terminal domain. Hierarchically coupled regulatory elements linking the NTD’s structural dynamics to channel activity were mapped along the entire length of the IDR. These elements modulate channel activity through lipid-dependent transient crosstalk. These results highlight important regulatory functions of the IDR and underscore that the IDRs cannot be neglected when trying to understand TRP channel structure and function.

## Results

### Structural ensemble of the TRPV4 N-terminal intrinsically disordered region

To address the current lack of structural and dynamic information for the TRPV4 NTD, we purified the 382 amino acid *Gallus gallus* domain (residues 2-382, with 83/90% sequence identity/similarity to human TRPV4) as well as its isolated IDR (residues 2-134), and ARD (residues 135-382) (Fig. 1a-c, Fig. S1). The avian proteins were chosen due to their increased stability compared to their human counterparts^27^. Analytical size-exclusion chromatography (SEC) and SEC-MALS (SEC multi-angle light scattering) showed that these constructs are monomeric, while circular dichroism (CD) spectroscopy and the narrow chemical shift dispersion of the [^1^H, ^15^N]-NMR (nuclear magnetic resonance) spectra of the ^15^N-labeled TRPV4 IDR in isolation or in the context of the NTD confirmed its high amount of disorder^5^ (Fig. 1c-e; Fig. S2).

**Figure 1:**
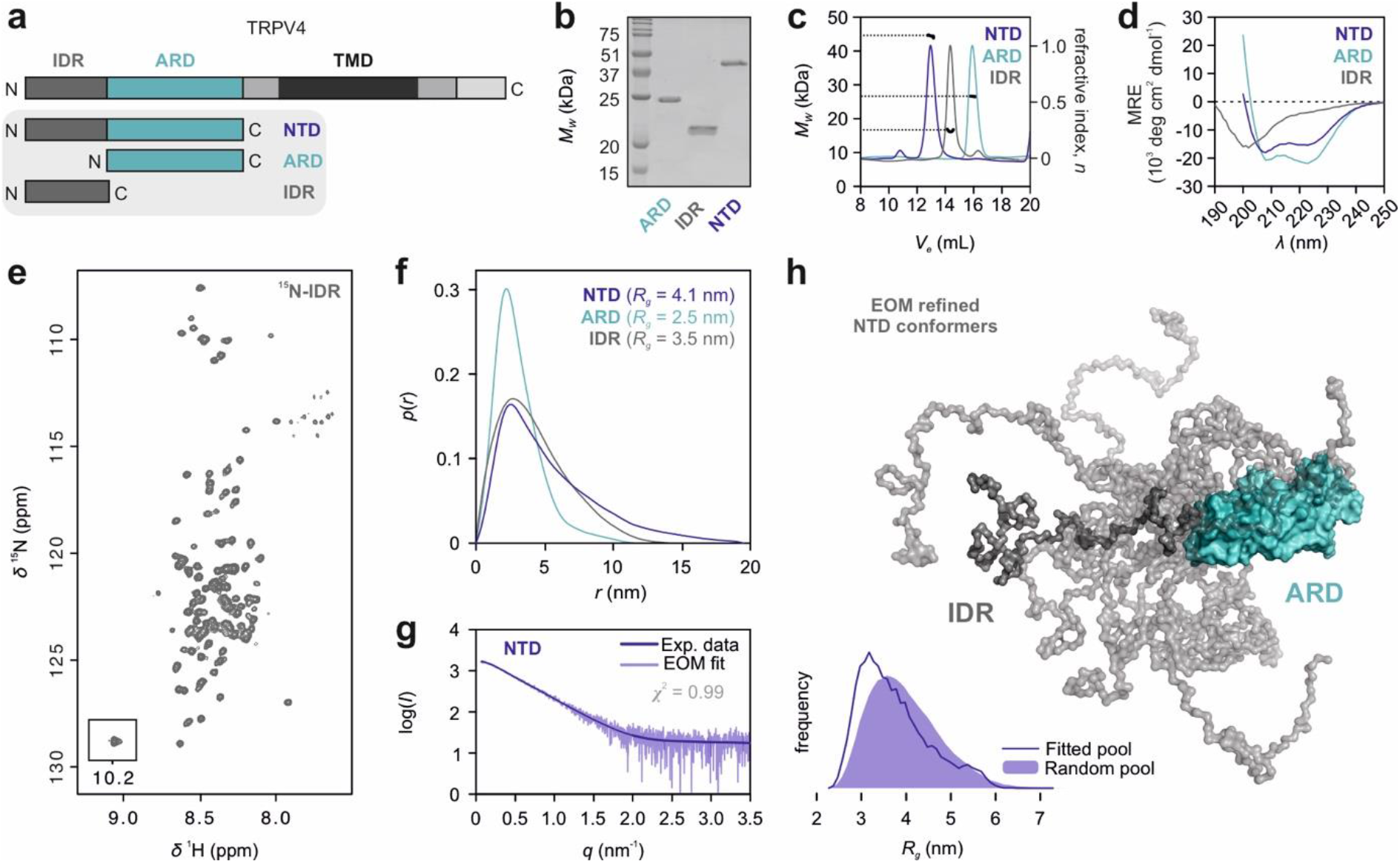
Structural ensemble of the TRPV4 N-terminal domain. **a** TRPV4 N-terminal constructs used for structural analyses. **b, c, d** Purified TRPV4 N-terminal constructs analyzed by Coomassie-stained SDS-PAGE (b), SEC-MALS (c) and CD spectroscopy (d). **e**[^1^H, ^15^N]-TROSY-HSQC NMR spectrum of ^15^N-labeled TRPV4-IDR (see Fig. S2 for backbone assignments). **f, g** SAXS pair-distance-distribution (f) and ensemble optimization method (EOM) analysis (g) of TRPV4 N-terminal constructs (see also Table S1 and Fig. S3). **h** NTD ensemble refined by EOM. A library of 10,000 NTD structures generated with a chain of dummy residues for the IDR and the X-ray structure of the TRPV4 ARD (PDB: 3W9G) as templates was refined against the experimental data. The fitted pool was compared to the random pool to select the sub-set of ensemble-states representing the experimental data. Ten IDR conformers best representing the experimental scattering profile are depicted.

An IDR-containing protein is best described as a structural ensemble, which can be analyzed by SEC-coupled small-angle X-ray scattering (SEC-SAXS) and subsequent Ensemble Optimization Method (EOM) analysis^34,35^. The isolated TRPV4 IDR is highly flexible and fluctuates between numerous conformations that, as a population, produce a skewed real-space scattering pair-distance distribution function, or *p*(*r*) profile that extends to **~** 12.5-15 nm (Fig. 1f, Fig. S3). Suggesting the presence of transient intradomain contacts, the TRPV4 IDR preferentially sampled more compact states both in isolation and attached to the ARD compared to a randomly generated pool of solvated, self-avoiding walk structures (Fig. 1f-h, Fig. S3). Interdomain contacts between the IDR and ARD were also apparent from the loss of IDR signal intensities in the ^1^H, ^15^N-NMR spectra of the isolated ^15^N-labeled IDR compared to the NTD (Fig. S2b, c), e.g. for residues ~20-35 and ~55 to 115. The ARD itself was not resolved in the spectra of the NTD likely due to unfavorable dynamics (see below).

### Structural dynamics of the TRPV4 ARD

The SAXS data, the dimensionless Kratky plot and the resulting *p*(*r*) profile of the isolated ARD are typical of a compact globular particle (Fig. S3). Accordingly, the 28 kDa ARD has a significantly smaller radius of gyration (*R_g_*~2.5 nm) and maximum particle dimension (*D_max_*~11.5 nm) than the 15 kDa IDR (*R_g_*~3.5 nm; *D_max_***~** 12.5-15 nm). Nonetheless, the SAXS data of the ARD could not be fitted with the scattering curves calculated from the available compact ARD X-ray crystal structures^10,17,18^ (Fig. S3f). Instead, models undergoing major conformational rearrangements had to be generated to obtain satisfactory fits to the experimental data of the ARD in solution. *Ab-initio* bead modeling using DAMMIN^36^ yielded a prolate-shape with a protrusion that may be consistent with the partial unfolding of one or two peripheral ankyrin repeats (Fig. 2a). Rigid-body normal-mode analysis of the ARD with SREFLEX^37^ suggested that a shift in the spatial disposition of the individual ankyrin repeats is required to satisfy the experimental data (Fig. 2b). As electron density in TRPV channel structures is frequently missing for the N-terminal ARD tips^4,38,39^, and melting temperatures of ~37 °C have been reported for the TRPV1 and TRPV4 ARDs^17,40^, TRPV channel ARDs may indeed fluctuate between structured and partially unstructured states.

**Figure 2:**
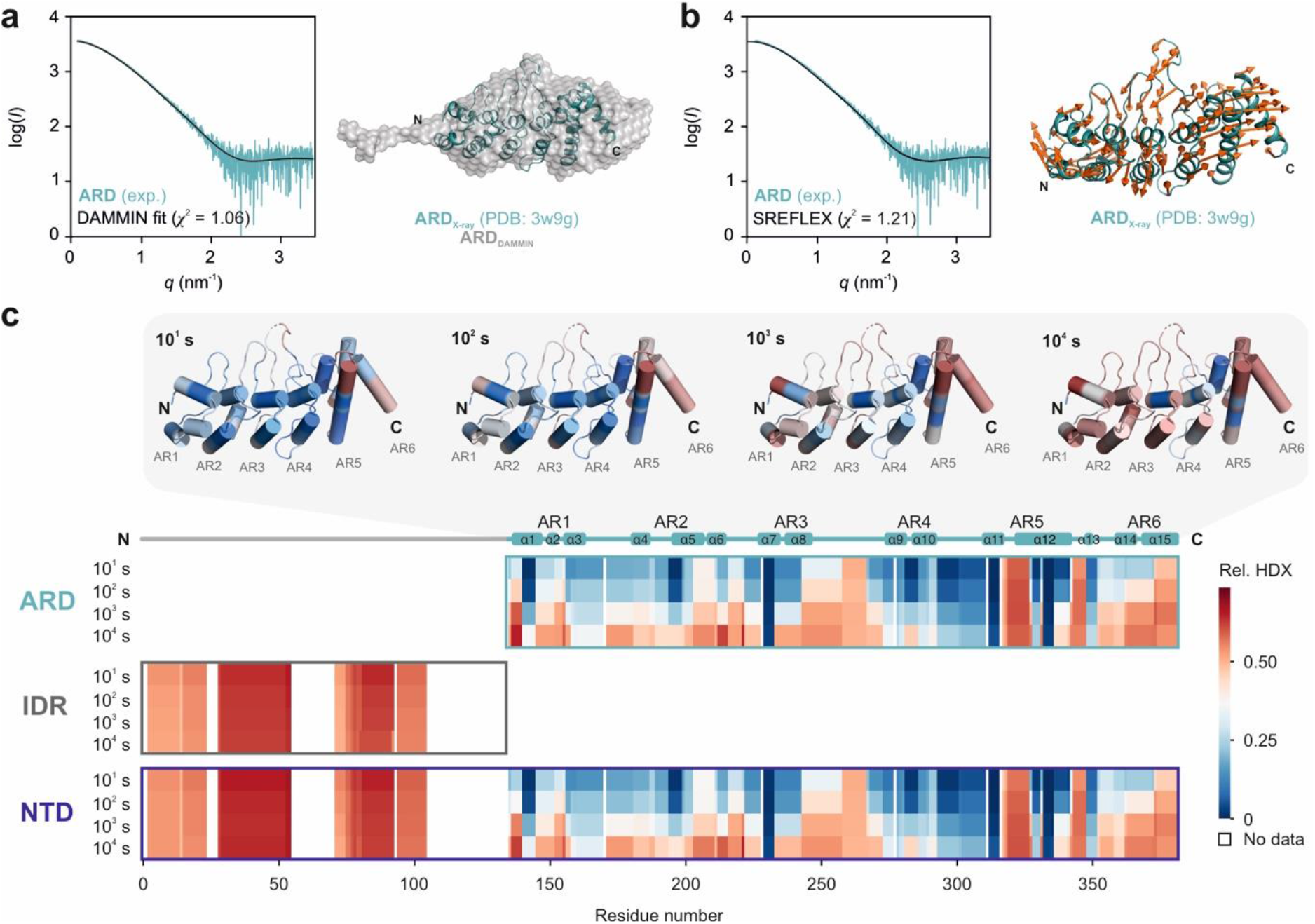
Structural dynamics of the TRPV4 ARD in solution. **a, b** For a better fit with the experimental SAXS data of the ARD in solution, DAMMIN (a) and SREFLEX (b) modeling of the TRPV4 ARD was carried out. The SAXS-based DAMMIN model (grey) is shown in comparison to the X-ray structure of the *G. gallus* TRPV4 ARD (PDB: 3W9G, teal). Normal mode vectors of the aligned SREFLEX model are shown with orange arrows and indicate which ARD regions undergo conformational rearrangements to satisfy the fits to the experimental data. **c** H/D exchange of TRPV4 NTD and its isolated subdomains. Low (blue) to high (red) HDX shown for four time points (see Supplemental Dataset 1). Areas without HDX assignment are colored white. For the ARD, HDX was visualized on the *G. gallus* TRPV4 ARD X-ray structure (PDB: 3W9G). Its topology with six ankyrin repeats (AR) is shown on top of the heat map diagram.

To further evaluate the structural flexibility of the ARD in solution, we used HDX-MS (hydrogen/deuterium exchange mass spectrometry) (Fig. 2c, Supplemental Data Set 1). HDX-MS probes the peptide bonds’ amide proton exchange kinetics with the solvent and thus provides insights into the higher order structure of proteins and their conformational dynamics^41^.

Immediate high HDX was apparent for ARD loop 3 (residues 259-267), ankyrin repeat 5 (residues 319-327) and the linker between ankyrin repeats 5 and 6 (residues 344-348). Most α-helices showed progression in HDX over time except for α7 (repeat 3), α9/α10 (repeat 4), and α11/α12 (repeat 5) suggesting that these constitute the structural core of the ARD with the least flexibility. The peripheral ankyrin repeats 1, 2 and 6 underwent faster exchange (HDX at 10^3^ s). The combined observations from HDX-MS and SAXS demonstrate that the ARD, although globally compact, experiences complex conformational dynamics, i.e., slower motions in the ARD core and faster dynamics within the ARD loops and peripheral ankyrin repeats in agreement with the extensive line broadening observed in the [^1^H, ^15^N]-NMR spectrum of the NTD (Fig. S2b).

For the IDR, immediate high HDX was apparent in the resolved parts (residues 3-24, 29-55, and 72-105) substantiating its unstructured character. The transient nature of interdomain contacts between ARD and IDR was underscored by the absence of a significant difference in the HDX of the individual domains in isolation or in the context of the full-length NTD.

### Long-range TRPV4 NTD interactions center on the PIP_2_-binding site

Long-range interactions between IDR and ARD were investigated using crosslinking mass spectrometry (XL-MS). Except for the first 49 amino acids, 25 lysine residues are almost evenly distributed throughout the *G. gallus* TRPV4 NTD sequence. The lysine side chain amino groups can be crosslinked by disuccinimidyl suberate (DSS), probing Cα-Cα distances up to 30 Å^42^. Both intradomain (within IDR or ARD) and interdomain (between IDR and ARD) crosslinks were observed for the NTD (Fig. 3a, Supplemental Data Set 2). Many intra- and interdomain contacts were observed for the most N-terminal IDR lysine residues (K50, K56) and those within or close to the PIP_2_-binding site on the C-terminal end of the IDR (K107, K116, K122). Importantly, these crosslinks were replicated in an equimolar mix of isolated IDR and ARD, supporting the involvement of these IDR regions in specific long-range interactions (Fig. 3b, c,).

**Fig. 3:**
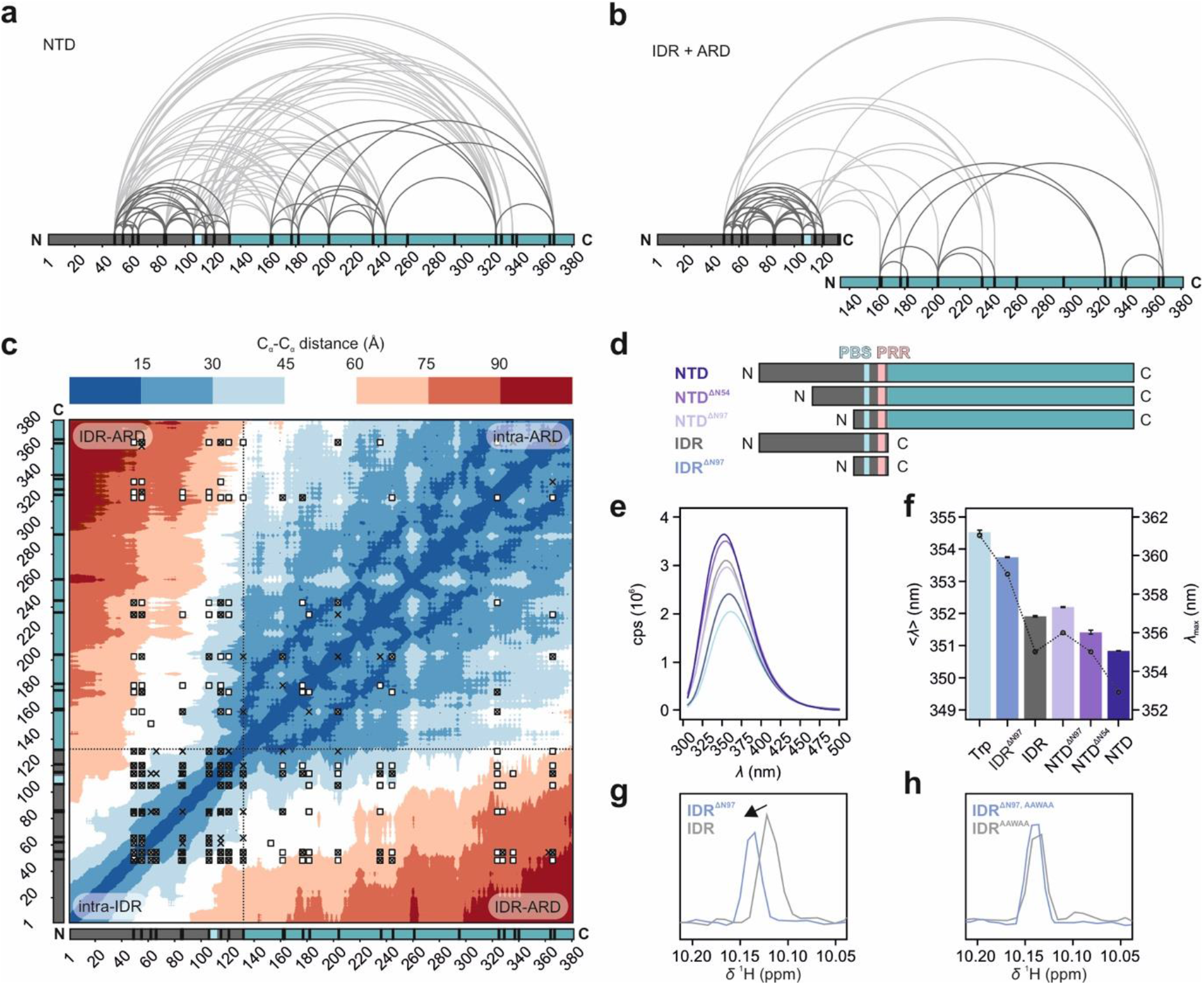
TRPV4 NTD long-range intra- and interdomain interactions. **a, b** Intra- and inter-domain interactions of IDR and ARD probed by crosslinking mass spectrometry. The entire NTD (a) or an equimolar mix of isolated IDR (grey) and ARD (cyan) (b) were used. Intradomain and interdomain crosslinks are shown by curved lines (dark and light grey, respectively), lysine residues (black ticks) and PIP_2_-binding site (light blue) are indicated. **c** Heat map of C_α_-C_α_ distances for an NTD conformational ensemble consisting of 15 EOM-refined conformers (Fig. 1h). Crosslinks are highlighted by white squares (NTD), black crosses (equimolar ARD:IDR mixture) or white squares filled with black crosses (both experimental set-ups). **d** TRPV4 N-terminal constructs used for tryptophan fluorescence (PBS: PIP_2_-binding site, PRR: proline rich region)**. e, f** Tryptophan fluorescence spectroscopy of TRPV4 N-terminal constructs (IDR, NTD, NTDΔN54 and NTDΔN97 lacking the first 54 or 97 amino acids, respectively, and IDR^ΔN97^ (comprising PIP_2_-binding site, surrounding basic residues and proline rich region) or isolated amino acid in buffer (Trp). Residue W109 in the PIP_2_-binding site is the sole tryptophan residue in the entire NTD. Bars represent the intensity weighted fluorescence emission wavelength <*λ*> (left axis). Error bars represent SD of mean of n=3. The fluorescence emission maximum *λ*_max_ is shown by black circles connect through dotted lines (right axis). **g, h**^1^H chemical shift differences of W109 sidechain amide between IDR and IDR^ΔN97^ as well as their respective counterparts harboring the PIP_2_-binding site (^107^KRWRR^111^) mutation to ^107^AAWAA^111^.

Conveniently, the TRPV4 PIP_2_-binding site (consensus sequence KRWRR) important for TRPV4 sensitization^31^ contains the sole tryptophan residue within the ~43 kDa NTD (W109 in our constructs). Changes in its chemical environment, e.g. through altered protein contacts, can thus be probed directly by differences in the tryptophan fluorescence spectra of deletion constructs generated around the PIP_2_-binding site (Fig. 3d).

The fluorescence emission of a minimal construct (IDR^ΔN97^, residues 97-134), which included the PIP_2_-binding site and the proline-rich region, was suggestive of high solvent accessibility of the tryptophan residue and resembled that of free tryptophan in buffer (Fig 3e, f). In longer constructs containing additional parts of the IDR, the ARD or both, the fluorescence emission was blue shifted, indicating that W109 was in a more buried, hydrophobic environment. This effect was most pronounced for the full-length NTD. Deletion of the N-terminal half of the IDR (NTD^ΔN54^) yielded an intermediate emission wavelength between full-length NTD and NTD^ΔN97^, a construct comprising only the ARD, proline-rich region, and PIP_2_-binding site. This indicates that the local PIP_2_-binding site environment is influenced by both the ARD and the distal IDR N-terminus.

### The PIP_2_-binding site promotes compact NTD conformations

To probe the role of the PIP_2_-binding site for the NTD conformational ensemble, we replaced its basic residues by alanine (KRWRR → AAWAA) across TRPV4 N-terminal constructs (Fig. S4a-d). CD spectroscopy and SEC showed that the structural integrity of the mutants was maintained (Fig. S4c, d). However, we noticed consistently higher Stokes radii compared to their native counterparts (Fig. S4e), suggesting that the charge neutralization of the PIP_2_-binding site affects the IDR structural ensemble. Likewise, the tryptophan emission wavelength of the AAWAA mutants was also increased compared to the native constructs, indicative of a more solvent-exposed central tryptophan residue (Fig. S4f, g). Furthermore, the ^1^H chemical shifts of the W109 sidechain amide were different between the native IDR and IDR^ΔN97^, but the same between the respective PIP_2_-binding site mutants (Fig. 3g, h). Thus, transient long-range interactions between the N-terminus and the PIP_2_-binding site seem to be disrupted upon mutation of the PIP_2_-binding site.

NTD^AAWAA^ and IDR^AAWAA^ were also analyzed by SEC-SAXS and EOM (Fig. 4, Fig. S4h-o). The scattering profile and real-space distribution of IDR^AAWAA^ resembled the native IDR, indicating a random chain-like protein. However, the mutant’s *R_g_* and *D_max_* values (3.5 nm and 14.5 nm, respectively) were slightly increased compared to the native IDR (*R_g_* = 3.4 nm and *D_max_* = 14.0 nm). This effect was even more pronounced in the context of the NTD, with *R_g_* and *D_max_* values of 4.5 nm and 19.5 nm, respectively, compared to *R_g_* = 4.1 nm and *D_max_* = 19.0 nm for the native IDR. Unlike the native IDR and NTD, the *R_g_* distributions of the mutant constructs agreed well with the randomly generated pools of solvated, self-avoiding walk structures (Fig. 4d). This suggests that constructs with a mutated PIP_2_-binding site populate expanded conformations more frequently and show more random chain-like characteristics, thereby substantiating the role of the PIP_2_-binding side as a central mediator of long-range contacts within the TRPV4 NTD.

**Fig. 4:**
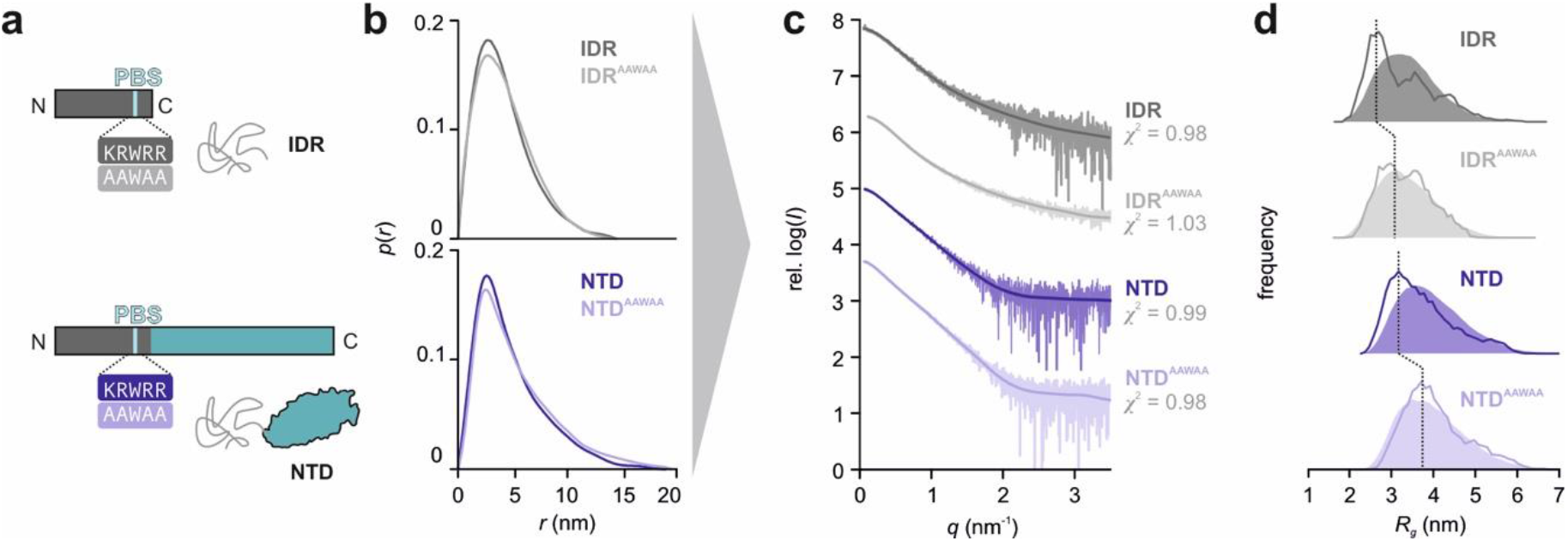
The PIP_2_-binding site promotes compact IDR conformations. **a** Constructs used in SEC-SAXS experiments. **b** Real-space pair-distance distribution function or *p*(*r*) profiles calculated for IDR and IDR^AAWAA^ (grey curves) as well as NTD and NTD^AAWAA^ (blue curves). *p*(r) functions were scaled to an area under the curve of 1. The real-space distance distribution of IDR^AAWAA^ yields a radius of gyration (*R_g_*) = 3.5 nm with a maximal particle dimension (*D_max_*) = 14.5 nm (native IDR: *R_g_* =3.4 nm, *D_max_* = 14.0 nm). NTD^AAWAA^ has a *R_g_* = 4.5 nm and a *D_max_* = 19.5 nm (native NTD: *R_g_* =4.1 nm, *D_max_* = 19.0 nm). **c** Fit between EOM-refined IDR and NTD models and experimental scattering data. **d** Comparison between *R_g_* values of IDR and NTD variants between random pool structure library (solid area) and EOM refined models (dotted line).

### Competing attractive and repulsive interactions between distinct IDR regions govern the NTD structural ensemble

The TRPV4 IDR consists of alternating highly conserved and non-conserved regions arranged along a charge gradient. An N-terminus rich in acidic residues segues into a C-terminus with an accumulation of basic residues followed by the proline-rich region connecting to the ARD (Fig. 5a, b). To probe the effects of differently charged and conserved IDR regions on the NTD structural ensemble, consecutive N-terminal deletion constructs were investigated by CD spectroscopy, SEC, and SEC-SAXS (Fig. 5c-e, Fig. S5, Fig. S6). The respective *R_S_*, *R_g_* and *D_max_* values for consecutive N-terminal deletions do not change linearly, rather, depending on their charge (*z*), individual IDR regions mold the structural ensemble of the NTD differently (Fig. 5e). Addition of only the proline-rich region to the ARD (NTD^ΔN120^) notably increased the *R_S_*, *R_g_* and *D_max_* values. This expansion is likely due to the formation of a polyproline helix^27^. A construct containing both the PIP_2_-binding site and the basic residues preceding it (NTD^ΔN97^) showed an increase in compaction over the construct with the PIP_2_-binding site alone (NTD^ΔN104^) suggesting cumulative effects of the regions surrounding the PIP_2_-binding site for NTD structure compaction. Adding another ~40 residues yields NTD^ΔN54^, which includes the entire basic and highly conserved central stretch of the IDR, did not significantly increase the protein dimensions further underscoring the importance of this region for interdomain crosstalk. This is supported by NMR spectroscopy, where the region between residues 55-115 showed notable peak broadening in the context of the entire NTD compared to the isolated IDR (Fig. S2c). Finally, the full-length NTD had significantly increased protein dimensions compared to NTD^ΔN54^, indicating that the overall structural ensemble of the TRPV4 NTD is modulated by competing attractive and repulsive influences exerted by distinct IDR regions.

**Fig. 5:**
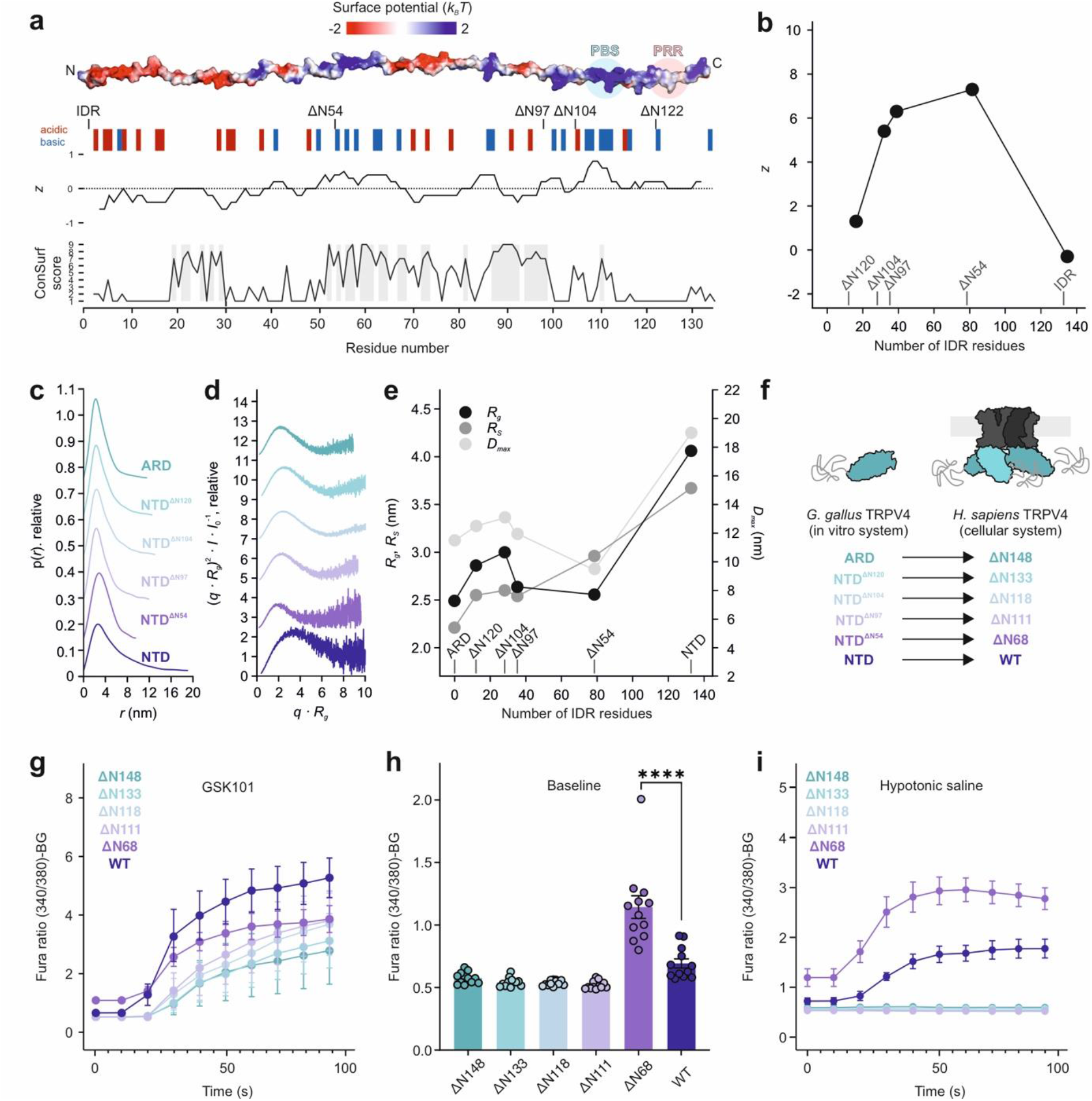
The distal IDR N-terminus affects the structural NTD ensemble and attenuates TRPV4 channel activity. **a** Topology of NTD truncations showing the charge distribution *z* and sequence conservation along the IDR. **b** Overall charge (*z*) at pH 7.4 of IDR deletion constructs. **c, d** Normalized real-space distance distribution *p*(*r*) and dimensionless Kratky plot of NTD and NTD deletion constructs. **e** Radius of gyration (*R_g_*) and Stokes radius (*R_S_*) determined from the real-space distance distribution (c) and SEC analysis (Fig. S5c), as well as maximum particle dimension (*D_max_*, right y-axis), plotted against number of IDR residues in NTD constructs. **f** N-terminal deletion mutants in the *in vitro* (*G. gallus*) and *in cellulo (H. sapiens)* systems. **g** Activation of hsTRPV4 constructs expressed in MN-1 cells with the specific agonist GSK101 at t=20 sec. **h** Basal Ca^2+^ levels in MN-1 cells expressing hsTRPV4 constructs. **i** Stimulation of Ca^2+^ flux by hypotonic saline at t=20 sec in MN-1 cells expressing different hsTRPV4 constructs. (n=12 wells with 10-30 cells/well for all Ca^2+^-influx experiments).

### The IDR N-terminus autoinhibits channel activity

To investigate the role of individual IDR regions on channel function, Ca^2+^ imaging of human TRPV4 N-terminal deletion constructs expressed in the mouse motor neuron cell line MN-1 was performed as described previously^29^ (Fig. 5f-i). All constructs were successfully targeted to the plasma membrane and structurally intact, as seen by the ability of the synthetic agonist ‘GSK101’^43^ to reliably activate the proteins (Fig. 5g, Fig. S7). All mutants had basal Ca^2+^ levels similar to the full-length channel. Only *H. sapiens* TRPV4^ΔN68^ (corresponding to *G. gallus* TRPV4^ΔN54^) had strongly increased basal Ca^2+^ levels (Fig. 5h). Deletion of the entire IDR (hsTRPV4^ΔN148^), as well as constructs retaining additional IDR regions, i.e., the proline-rich region (hsTRPV4^ΔN133^/ggTRPV4^ΔN120^), the PIP_2_-binding site (hsTRPV4^ΔN118^/ggTRPV4^ΔN104^) and the preceding basic residues (hsTRPV4^ΔN111^/ggTRPV4^ΔN97^) yielded a channel non-excitable for osmotic stimuli (Fig. 5i). In contrast, hsTRPV4^ΔN68^ was hypersensitive to osmotic stimuli and its Ca^2+^ influx far exceeded that of the native channel, indicating that the IDR N-terminus acts as a dominant autoinhibitory element. Furthermore, the data show that the PIP_2_-binding site is not sufficient for osmotic channel activation but additionally requires the presence of the central IDR around residues ~68-111 (~54-97 in ggTRV4).

### The IDR N-terminus attenuates IDR lipid binding

Lipids and lipid-like molecules are important TRPV4 functional regulators^31,44–46^, but beyond the PIP_2_-binding site, lipid interactions with the TRPV4 NTD have not been probed in detail. A previously proposed lipid binding site in the ARD^44^ seems implausible because it does not face the membrane in the context of the full-length channel^19,20^. Indeed, neither the isolated ARD, nor NTD^ΔN120^, also containing the proline-rich region, interacted with POPC/POPG liposomes in a sedimentation assay (Fig. 6a, b, Fig. S8a, b). In contrast, ~75% of the full-length NTD was found bound to liposomes. For NTD^AAWAA^, lipid binding was reduced to ~20%, indicating that the PIP_2_-binding site is a major, but not the only lipid interaction site in the TRPV4 IDR. Deletion of the IDR N-terminal half slightly increased the fraction of lipid-bound protein. Incidentally, “protection” of the PIP_2_-binding site from lipid binding by the N-terminal IDR was also observed with tryptophan fluorescence (Fig. S8c-g) and may indicate that long-range intra-domain contacts compete with lipid binding in the native IDR.

**Fig. 6:**
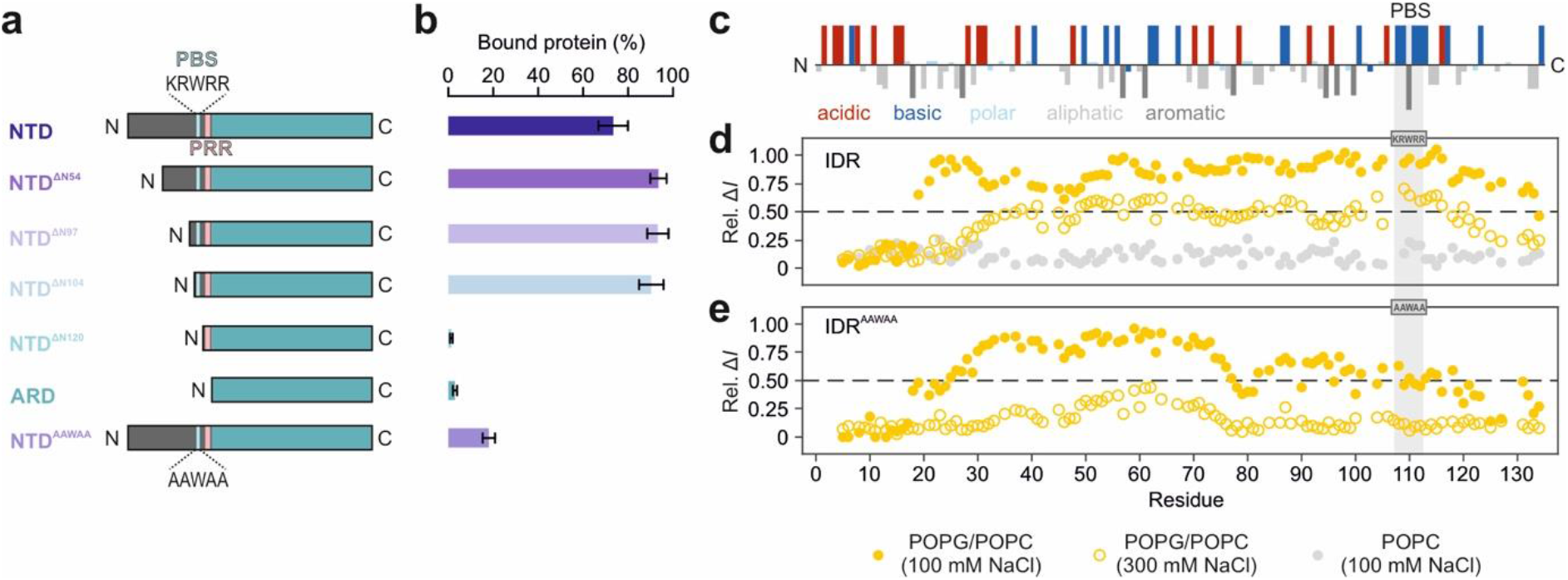
The TRPV4 IDR interacts extensively with lipids. **a** Topology of N-terminal deletion mutants used for liposome sedimentation assay. **b** Protein distribution between pellet (’bound protein’) and supernatant after centrifugation, quantified via gel densitometry. Error bars represent SD of mean from n=3. **c** TRPV4 IDR residues are arranged along a charge gradient. **d, e** NMR signal intensity differences for ^15^N-labeled IDR variants (100 μM) titrated with POPC (grey circles) or POPC/POPG liposomes at low (yellow circles) or high salt concentration (empty circles). Higher values are indicative of lipid binding.

NMR chemical shift perturbation assays allowed identification of the lipid-interacting IDR residues (Fig. 6c-e, see Fig. S9 for ^13^C, ^15^N-labeled IDR^AAWAA^ backbone assignments). In the native IDR, ~75% of all residues showed line-broadening in the presence of POPG-containing liposomes. Coarse-grained molecular dynamics (MD) simulations of the IDR on a plasma membrane mimetic corroborated the NMR experiments (Fig. S10, Table S3). Lipid interactions were seen to be dominated by the PIP_2_-binding site, the central IDR and a conserved N-terminal patch (see below). In MD simulations of IDR^AAWAA^, lipid interactions were severely reduced in the PIP_2_-binding site, again agreeing with the NMR data (Fig. 6e, Fig. S10).

Indicating an electrostatic contribution, an increase in salt concentration or the use of net-neutral POPC liposomes in NMR experiments reduced the observed lipid-induced line broadening for both native IDR and IDR^AAWAA^ (Fig. 6d, e). Interestingly, the MD simulations also suggested a general preference for negatively charged PIP_2_ over other membrane constituents for both the PIP_2_-binding site and the central IDR (Fig. S10b, c).

Since we saw that N-terminal deletion mutants retaining the PIP_2_-binding site essential for channel sensitization^31,46^ are still inactive if they do not also include the central IDR (Fig. 5i), the function of the central IDR may be two-fold – enriching PIP_2_ in the channel vicinity and increasing the IDR’s residency time at the plasma membrane.

### A conserved patch in the IDR N-terminus mediates transient long-range interactions and autoinhibits TRPV4

We hypothesized that the site(s) in the N-terminal half of the IDR responsible for the observed channel autoinhibition and lipid binding attenuation may act via the PIP_2_-binding site. Thus, we compared the native IDR and IDR^AAWAA^ NMR backbone amide chemical shifts to reveal interactions between the N- and C-terminal ends of the IDR (Fig. 7a). The largest chemical shift differences were naturally found in and around the mutated PIP_2_-binding site itself and to a lower degree in the central IDR. Additionally, a region encompassing residues ~20-30 in the IDR N-terminus also showed notable chemical shift differences. This patch is the only conserved stretch in the N-terminal half of the IDR (consensus sequence FPLS-S/E-L-A/S-NLFE (^19/31^FPLSSLANLFE^29/41^ in gg/hsTRPV4)) (Fig. 7b, Fig. S11).

**Fig. 7:**
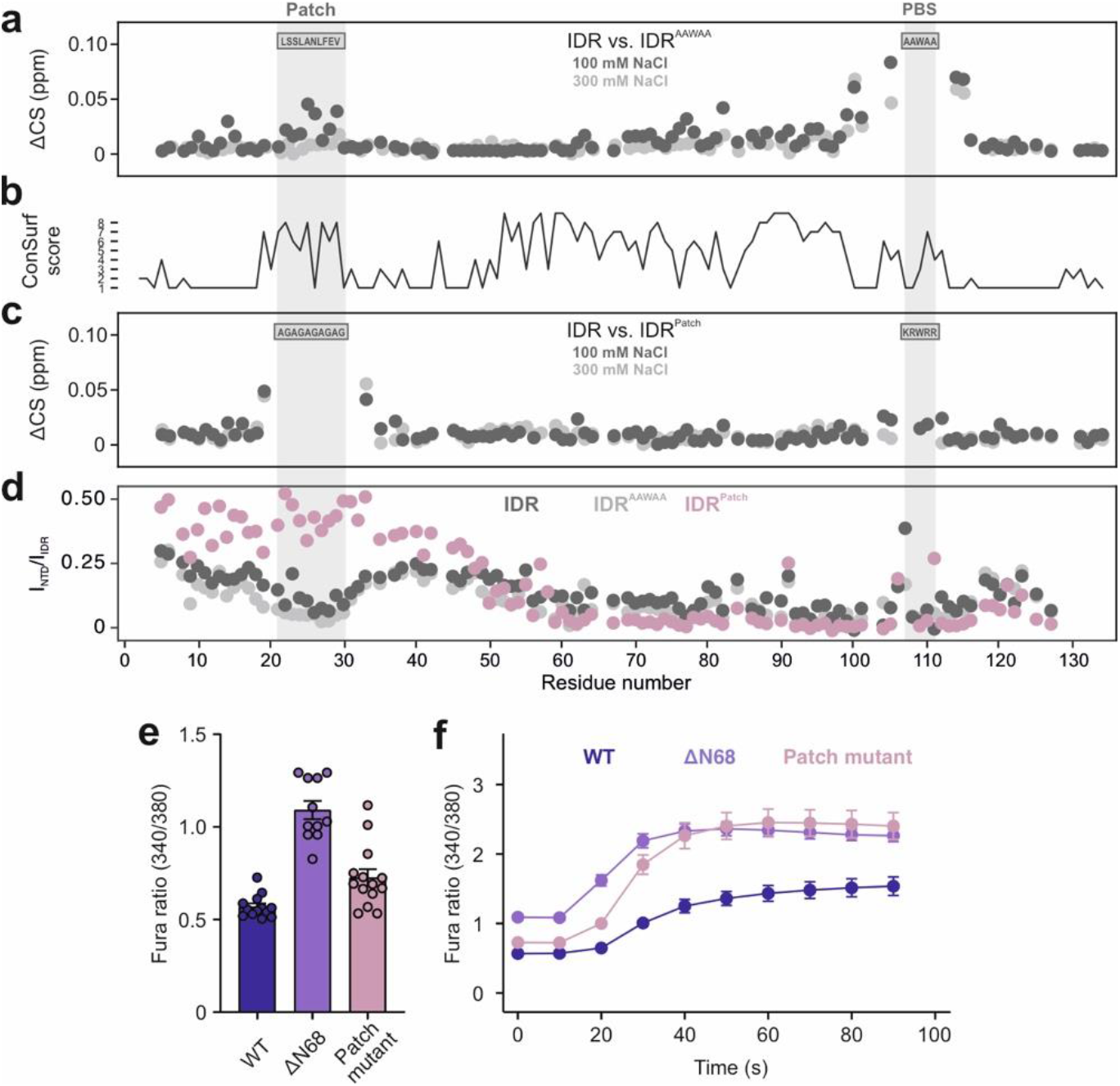
An N-terminal conserved patch transiently interacts with the PIP_2_-binding site and autoinhibits TRPV4. **a** Comparison of chemical shifts between ^15^N-labeled IDR and IDR^AAWAA^. A PIP_2_-binding site (PBS) mutation also affects the highly conserved N-terminal patch. At higher salt concentrations (light grey), these interactions are significantly reduced. **b** Degree of TRPV4 IDR conservation (compare Fig. S11). **c** Mutation of the N-terminal patch (IDR^Patch^) also affects the PIP_2_-binding site. **d** Relative peak intensities between isolated IDR and their corresponding NTD constructs (at 100 μM). Values <1 are indicative of IDR/ARD interactions, a value of zero represents complete line broadening of IDR resonances in the context of the NTD. **e, f** Ca^2+^ imaging of hsTRPV4 variants expressed in MN-1 cells. (e) Basal Ca^2+^ (n=13 (TRPV4), 11 (TRPV4^ΔN68^), 14 (TRPV4^Patch^)) and (f) hypotonic treatment at t = 20 sec (n=12 wells with 10-30 cells/well) show increased activity of the patch mutant. For better comparison, data for TRPV4^ΔN68^ are replotted from Fig. 5h, i.

Since our NMR data showed that the patch region is unstructured^5^ (Fig. S2), we replaced it with an (AG)_5_ repeat to avoid α-helix formation (IDR^Patch^, Fig. S11, see Fig. S12a, b for backbone NMR assignment). NMR relaxation data confirmed the absence of transient structure formation in the IDR^Patch^ mutant (Fig. S12c, d). Importantly, the resonances of the IDR^Patch^ PIP_2_-binding site residues showed chemical shift changes compared to the native IDR (Fig. 7c), suggesting that patch and PIP_2_-binding site on opposite ends of the IDR are in transient contact. Of note, the absence of persistent long-range interactions between the N-terminal patch and the PIP_2_-binding site for the IDR in the MD simulations (Fig. S10d) can be explained if the interactions are dominated by long-range electrostatics, which are comparably poorly represented in the coarse-grained simulation model. The patch apparently also undergoes additional transient interactions with the ARD since the NMR signal intensities of residues in and around the patch region in the native IDR and IDR^AAWAA^ showed significant line broadening within the context of the NTD but not in the isolated IDR. This effect was abrogated in the IDR^Patch^ mutant (Fig. 7d).

To elucidate the role of the patch for channel function, the full-length human TRPV4 channel harboring the patch mutant was expressed in MN-1 cells (Fig. 7e, f, Fig. S7b, Fig. S11). Compared to the native channel, TRPV4^Patch^ displayed significantly increased basal Ca^2+^ levels and osmotic hyperexcitability, as previously seen for TRPV4^ΔN68^. This shows that the conserved patch in the N-terminal IDR is the dominant module responsible for autoinhibiting channel activity.

### The conserved patch competes with PIP_2_ for binding to the PIP_2_-binding site

To probe whether the autoinhibitory patch also influences PIP_2_ binding to the IDR, we carried out NMR chemical shift perturbation assays (Fig. 8a-c). In the native IDR, residues within and around the PIP_2_-binding site (residues ~100-115), the central IDR (residues ~55-100) and the autoinhibitory patch (residues ~20-30) show the strongest responses to diC8-PIP_2_ addition (Fig. 8a). In our coarse-grained simulations, a substantial local increase in PIP_2_ was observed around the PIP_2_-binding site and the central IDR, but not in the patch region (Fig. S10b, c). This indicates that the observed NMR chemical shifts within the N-terminal patch are secondary effects, presumably based on altered protein-protein interactions upon PIP_2_ addition. Notably, in the native IDR, both PIP_2_-binding site and patch showed a similar dose response to PIP_2_ as gauged by the similar degree of line broadening for these regions (Fig. 8a, grey bars). This suggests that PIP_2_-binding site and patch act in concert and that lipid interactions in the PIP_2_-binding site are also sensed by the autoinhibitory patch.

**Fig. 8:**
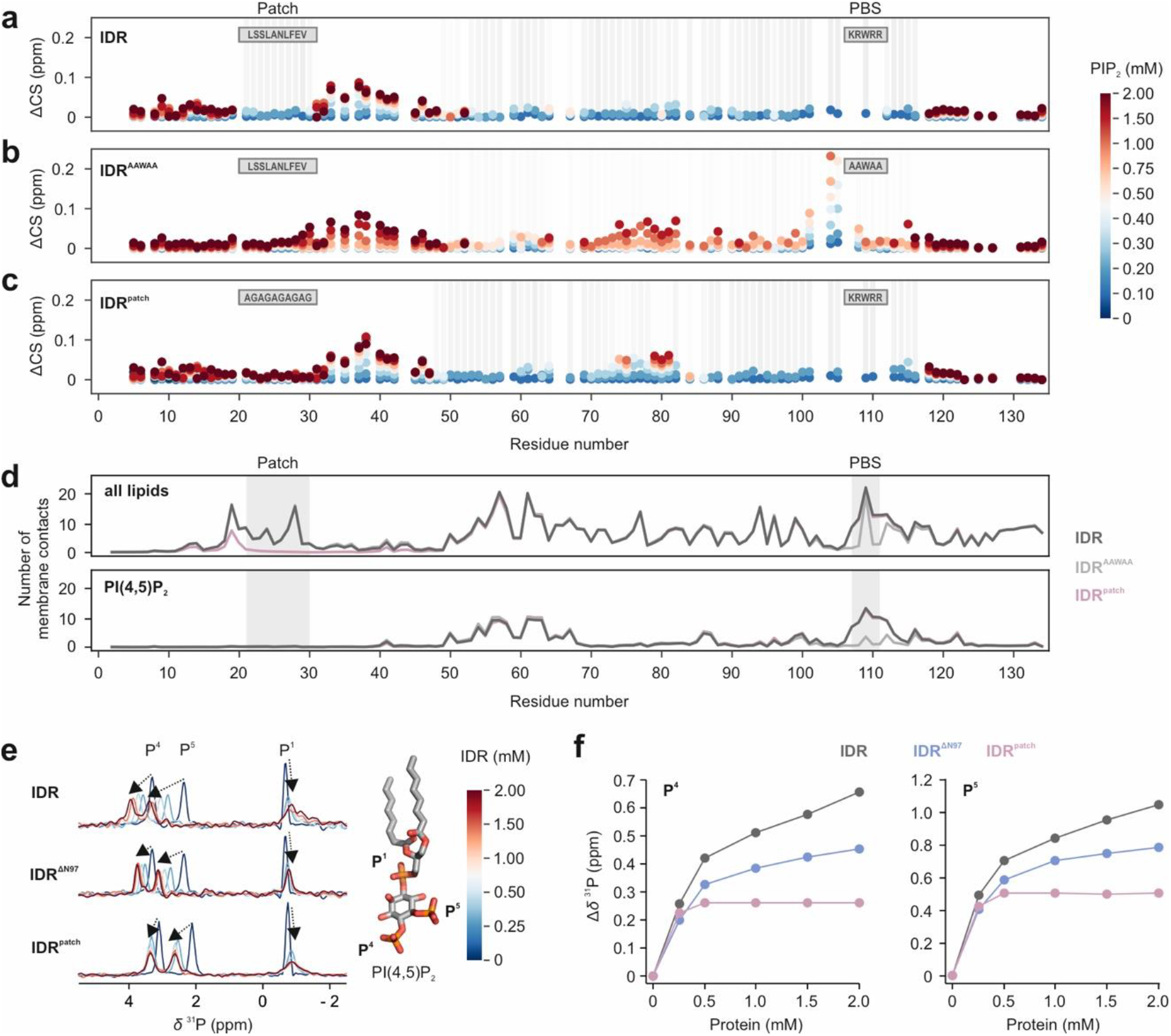
The N-terminal patch modulates PIP_2_ binding to the IDR. **a, b, c** Chemical shift perturbation of ^15^N-labeled IDR (a), IDR^AAWAA^ (b) and IDR^Patch^ (c) titrated with diC_8_-PIP_2_. Chemical shift changes are depicted by colored spheres, line broadening is indicated by grey bars. **d** Average number of membrane contacts from coarse-grain MD simulations with the native IDR (dark grey), IDR^AAWAA^ (light grey) or IDR^Patch^ (mauve) on a lipid bilayer composed of POPC (69%), cholesterol (20%), DOPS (10%) and PIP_2_ (1%). Contacts with all lipids (top) and only PIP_2_ (bottom) are shown. Four replicate 38 μs simulations were carried out per construct, contact averages were calculated from the last ~28 μs of each simulation. **e**^31^P NMR spectra of diC_8_-PIP_2_ (light blue) titrated with IDR, IDR^DN97^ or IDR^Patch^. Chemical shift changes are indicated by arrows. **f** Extent of chemical shift perturbations of P4 and P5 lipid headgroup resonances upon addition of IDR (grey), IDR^ΔN97^ (blue) or IDR^Patch^ (mauve).

For IDR^AAWAA^, dampened spectral responses to PIP_2_ were observed in the mutated PIP_2_-binding site, large parts of the central IDR and, to a much lesser degree, in the patch region (Fig. 8b) suggesting reduced coupling between these regions when the PIP_2_-binding site is mutated. Likewise, in MD simulations of IDR^AAWAA^, PIP_2_ binding was largely abrogated in the mutated PIP_2_-binding site (Fig. 8d, Fig. S10c). Consequently, the mutated PIP_2_-binding site frequently lost and regained contact with the lipid bilayer, although other IDR regions remained attached to the membrane throughout the simulations.

Mutation of the patch did not alter the lipid interaction pattern with the central IDR and PIP_2_-binding site *per se* as gauged by ^1^H, ^15^N-NMR spectroscopy and MD simulations (Fig. 8c, Fig. S10b, c, Fig. S12e). Since the severe line broadening in the ^1^H, ^15^N-NMR spectra precluded a more detailed analysis, we also took advantage of the PIP_2_ headgroup phosphate groups as a ^31^P NMR reporter (Fig. 8e). In agreement with the liposome sedimentation assay (Fig. 6), both the deletion (IDR^ΔN97^) or mutation of the patch (IDR^Patch^) mutation increased PIP_2_ binding compared to the native IDR as gauged by the extent of the respective ^31^P chemical shifts (Fig. 8f). Thus, the N-terminal patch seems to compete with PIP_2_ lipids for the PIP_2_-binding site via transient protein-protein interactions, thereby suppressing channel activity.

### Membrane-bound PIP_2_-binding site exerts a pull force on the ARD

It remains unclear how lipid binding to the IDR is transduced to the structured core of TRPV4 to modulate the conductive channel properties. In coarse-grained MD simulations, we emulated the positioning of an ARD-anchored IDR in a full-length channel by keeping the IDR’s C-terminal residue (V134) at distances of 5-9 nm from the membrane midplane (Fig. 9a). From the mean restraint forces for native IDR, IDR^AAWAA^ and IDR^Patch^, we determined force-displacement curves as a function of the distance between V134 and the membrane center (Fig. 9b).

**Figure 9:**
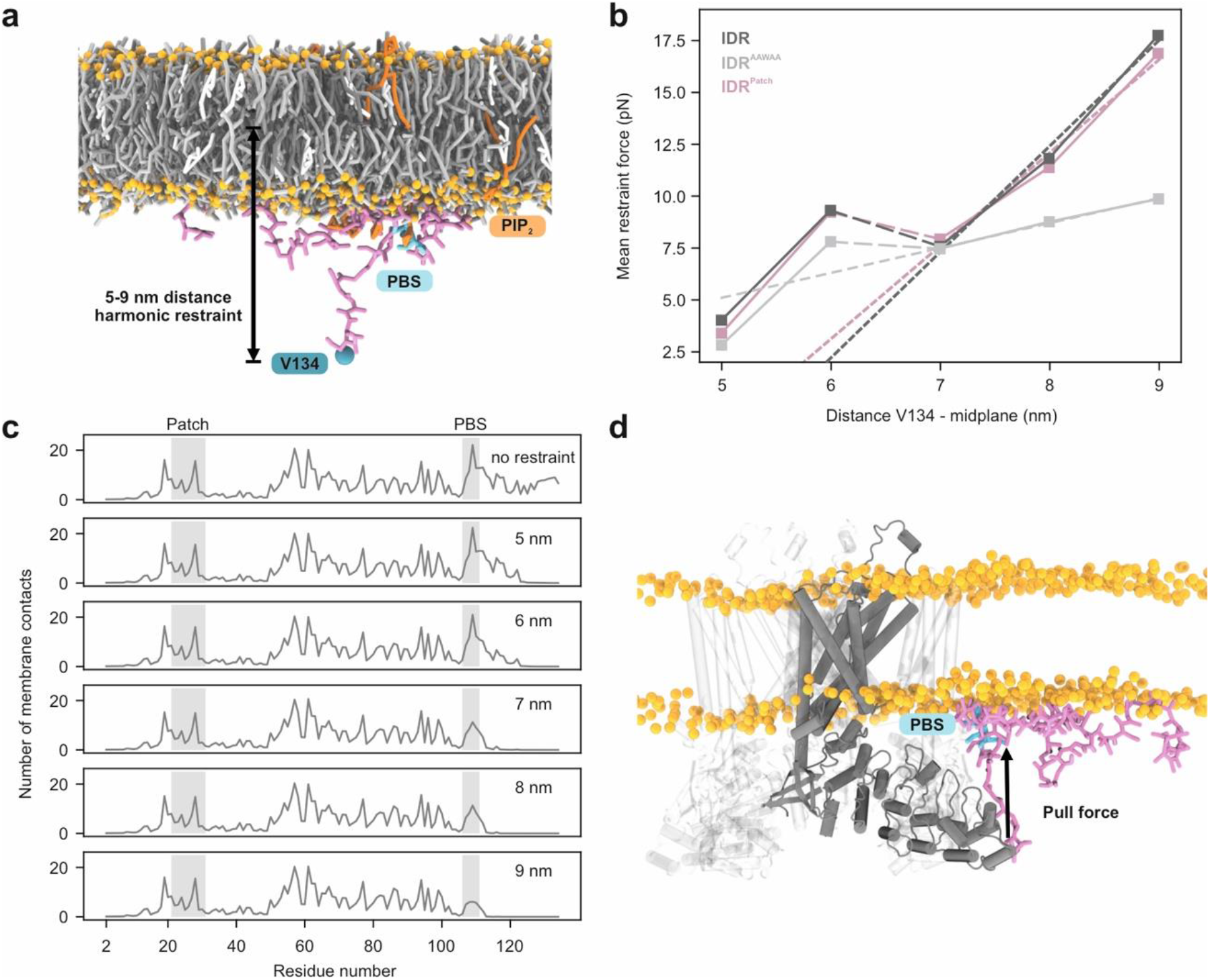
PIP_2_ binding to the TRPV4 IDR exerts a pulling force on the ARD. **a** Coarse-grained MD simulation of the TRPV4 IDR (pink liquorice) on a lipid bilayer containing 1% PIP_2_ (see Table S3). Headgroup phosphates are shown as orange spheres. The IDR C-terminus (V134; blue sphere) was held at defined distances from the membrane midplane to emulate anchoring by the ARD (PBS: PIP_2_-binding site, cyan). **b** Force-displacement curves from restrained simulations of TRPV4 IDR, IDR^AAWAA^ and IDR^Patch^. The mean restraint force is plotted against the mean distance between residue V134 and the membrane midplane. Dotted lines show linear fits to the regime dominated by PIP_2_-binding (>6.5 nm distance). Averages were calculated from the last ~28 μs of four 38-μs replicate simulations per construct and height restraint. Error bars present standard errors of the mean (SEM) of the replicate simulations, they are on the order 10^−3^ and thus not visible in the graph. **c** Membrane lipid contacts for each IDR residue at a given height restraint (see Fig. S10c for IDR^AAWAA^ and IDR^Patch^). Averages were calculated from the last 28 μs of each of the four replicate simulations. **d** Composite figure of a structure of the native IDR (from an MD simulation at a restraint distance of 7 nm) and an AlphaFold multimer model of the transmembrane core of the *G. gallus* TRPV4 tetramer. The force-displacement curves in (b) indicate that the interaction of the PIP_2_-binding site with the membrane exerts a pull force on the ARD N-terminus (solid arrow).

At heights <6.5 nm, all constructs experienced similar forces. At a height of ~6.5 nm, the residues C-terminal of the PIP_2_-binding site detached from the membrane in all IDR constructs. Thus, forces at heights ≥7 nm are generated by the PIP_2_-binding site pulling at the membrane. Importantly, the PIP_2_-binding site remained membrane-bound over the entire height regime in the simulations with native IDR and IDR^Patch^. In contrast, these interactions were lost in the IDR^AAWAA^ mutant, resulting in greatly reduced pull forces (~10 versus ~17.5 pN at 9 nm) (Fig. 9b, c). Furthermore, the reduced slope of the near-linear force-height curve beyond 6.5 nm implies a four-fold higher effective force constant acting on the IDR C-terminus with an intact PIP_2_-binding site compared to IDR^AAWAA^.

The forces observed here for the membrane-bound IDR are in the regime reported for other biochemical processes^47,48^. However, the smoothened energy landscape in our coarse-grained simulations may underestimate the actual force exerted on the ARD by its IDR “lipid anchor”. The strength of the PIP_2_-binding site interaction with the membrane also became apparent when constraining the IDR C-terminus at heights >8 nm. Here, rather than detaching, the pull of the PIP_2_-binding site led to noticeable membrane deformations (Fig. S10a). TRPV4 may thus not only be able to sense, but under certain conditions also directly affect its membrane microenvironment via its IDR.

### An integrated structural model of the TRPV4 N-terminal ‘belt’

Structural information for TRP channel IDRs is incomplete at best since they are not amenable to X-ray crystallography or cryo-electron microscopy studies due to their inherent spatiotemporal flexibility^4^. We previously calculated the dimensions theoretically sampled by TRP channel IDRs assuming they behaved as unrestrained worm-like chains and found that fully expanded IDRs of TRP vanilloid channels may contribute an additional 5–7 nm end-to-end distance to the structured cytosolic domains^4^. Here, by integrating our SAXS- and MD-derived IDR conformers into the structured TRPV4 core, we found that the cytosolic “belt” formed by the TRPV4 N-terminal IDRs is smaller due to their extensive lipid and intradomain interactions (Fig. 10a, b, Fig S13, Supplemental Movies S1 and S2). Nonetheless, the N-terminal IDRs more than double the TRPV4 diameter along the membrane plane from approximately 140 Å to a maximum of ~340 Å. With their IDRs, these proteins thus dramatically extend their reach and may act as multivalent cellular recruitments hubs.

**Fig. 10:**
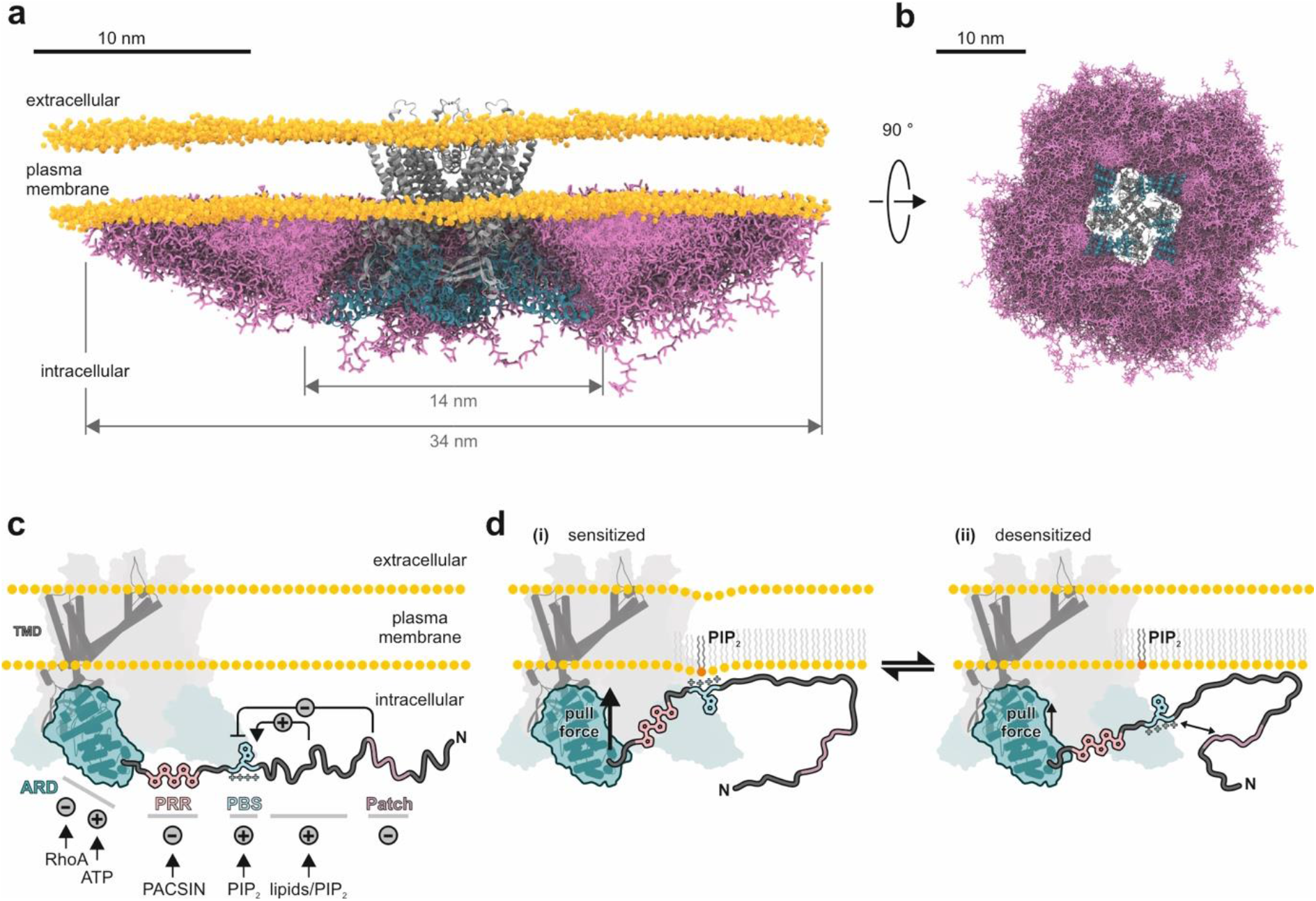
The N-terminal TRPV4 IDR significantly expands the protein dimensions and encodes a hierarchy of antagonistic regulatory modules. **a, b** Inclusion of the TRPV4 N-terminal IDR (pink licorice, ensemble from coarse-grained MD simulations and SAXS) more than doubles the dimensions of the full-length TRPV4 tetramer (AlphaFold multimer prediction of *G. gallus* TRPV4 transmembrane core (grey) and ARDs (cyan)) along the membrane plane as seen from the side (a) and the cytoplasm (b). 4012 unique IDR conformations are shown. For better visualization, the IDR conformations of the front facing TRPV4 monomer have been omitted in (a). Note that the distribution of the IDR conformers is not random, but rather governed by an intricate network of intra-domain and lipid interactions. For visualization of the IDR conformations within a single TRPV4 subunit, see Fig. S13 and Supplemental Movies S1, S2. **c** The TRPV4 N-terminus encodes antagonistic elements that regulate TRPV4 activity through ligand, protein, lipid or transient intra-domain contacts. **d** The membrane-bound PIP_2_-binding site exerts a pull force on the IDR C-terminus, presumably keeping TRPV4 in a sensitized state. The autoinhibitory patch modulates PIP_2_ binding and thus IDR membrane interactions, thereby attenuating channel activity.

## Discussion

In this study, we have shown that the TRPV4 N-terminal IDR encodes a network of transiently coupled regulatory elements that engage in hierarchical long-range crosstalk and can enhance or suppress TRPV4 activity (Fig. 10c). Such contacts may affect lipid binding as seen here, but presumably can also be modulated by other ligands^49^, regulatory proteins^27–30,49^ or post-translational modifications^50^ within the ARD and IDR to enable a fine-tuned integration of multi-parameter inputs by TRPV4.

The ARD is connected to the PIP_2_-binding site via a proline-rich region which forms a poly-proline helix^27^. The proline-rich region may thus be a relatively stiff connector to efficiently transduce pull forces between ARD and membrane-bound PIP_2_-binding site as suggested by our MD simulations (Fig. 9). Our NMR experiments show that the proline-rich region is not affected by lipids itself (Fig. 6d, 8a), but it binds the channel desensitizer PACSIN3^27,28,30^, which may affect the interaction between membrane-bound IDR and structured channel core. Likewise, the N-terminal autoinhibitory patch may reduce the pull force exerted by the PIP_2_-binding site by competing with its ability to bind lipids and thus effectively dampen channel activity. Our data thus provide a mechanistic explanation for prior observations that TRPV4 variants lacking part of the distal N-terminus display osmotic hypersensitivity^26,51^. Furthermore, a conformational equilibrium between PIP_2_-binding site interaction between membrane and autoinhibitory patch may allow TRPV4 to fine-tune channel responses depending on cell state and regulatory partners (Fig. 10d).

In summary, to understand TRP channel function and structure, their often extensive IDRs cannot be ignored. Our work shows that “IDR cartography”, i.e., mapping structural and functional properties onto distinct IDR regions through an integrated structural biology approach, can shed light on the complex regulation of a membrane receptor through its hitherto mostly neglected regions.

## Methods

### Antibodies and reagents

All chemicals were purchased from Sigma-Aldrich, Roth and VWR unless otherwise stated. Reagents used include GSK1016790A (‘GSK101’, Sigma-Aldrich, G0798), AlexaFluor 555 Phalloidin (ThermoFisher Scientific), ^15^N-NH_4_Cl and ^13^C_6_-glucose (EurisoTop). DSS-H12/D12 for crosslinking was obtained from Creative Molecules Inc. Lipids were purchased from Avanti Polar Lipids and Cayman Chemicals. Antibodies used were rabbit anti-GFP (Thermo Fisher Scientific, A-11122), rabbit anti-β-actin (Cell Signaling Technology, 4967) and HRP-conjugated monoclonal mouse anti-rabbit IgG, light chain specific (Jackson ImmunoResearch, 211-032-171).

### Computational Tools

Freely available computational tools were used to investigate the properties of N-terminal TRPV4 constructs. Sequence conservation was determined with ConSurf^52^ (Fig. 5, 7 and S11). Overall charge (*z*) and charge distribution of IDR deletion constructs were determined with ProtPi (www.protpi.ch and www.bioinformatics.nl/cgi-bin/emboss/charge) (Fig. 5). Gel densitometry analysis was carried out with ImageJ^53^ (Fig. 6, Fig. S8). The IDR charge gradient in Fig. 6c was plotted with the PepCalc tool (https://pepcalc.com/).

### Cloning, expression and purification of recombinant proteins

The DNA sequences encoding for the *G. gallus* TRPV4 N-terminal domain were cloned into a pET11a vector with an N-terminal His_6_SUMO-tag as described previously^27^. Human TRPV4 constructs in a pcDNA3.1 vector were commercially obtained from GenScript. Expression plasmids encoding for the isolated intrinsically disordered region (IDR), the isolated ankyrin repeat domain (ARD), N-terminal truncations (NTD^ΔN54^, NTD^ΔN97^, NTD^ΔN104^, and NTD^ΔN120^) and a peptide comprising residues 97-134 (IDR^Δ97^) were obtained from the NTD encoding vectors using a Gibson Deletion protocol^54^. Site-directed mutagenesis of the PIP_2_-binding site (^107^KRWRR^111^ to ^107^AAWAA^111^, forward primer GTGAAAACGCAGCCTGGGCCGCGCGTGTGGTTGAAAAACCAGTGG; reverse primer CACACGCGCGGCCCAGGCTGCGTTTTCACCACCAATCTGT) and regulatory patch (^19^FPLSSLANLFE^29^ to ^19^FP(AG)5E^29^, forward primer GATGACTCCTTCCCGGCCGGCGCGGGCGCCGGCGCGGGTGCGGGTGAGGACACCCCGTCT; reverse primer CGGGAAGGAGTCATCCCCCAGCACGTCCCC) were introduced in the abovementioned constructs by site-directed mutagenesis using polymerase chain reaction.

TRPV4 N-terminal constructs were expressed in *Escherichia coli* BL21-Gold(DE3) (Agilent Technologies) grown in terrific broth (TB) medium (or LB medium for IDR^Δ97^) supplemented with 0.04% (w/v) glucose and 0.1 mg/mL ampicillin. Cells were grown to an OD_600_ of 0.8 for induction with 0.5 mM IPTG (final concentration) and then further grown at 37 °C for 3 hrs. ^15^N, ^13^C-labeled proteins were prepared by growing cells in M9 minimal medium^55^ with ^15^N-HN_4_Cl and ^13^C-glucose as the sole nitrogen and carbon sources. Cells were grown at 37 °C under vigorous shaking to an OD_600_ of 0.4, moved to RT, grown to OD_600_ of 0.8 for induction of protein expression with 0.15 mM IPTG (final concentration) and then grown overnight at 20 °C. After harvest by centrifugation, cells were stored at −80 °C until further use. All purification steps were carried out at 4 °C. Cell pellets were dissolved in lysis buffer (20 mM Tris pH 8, 20 mM imidazole, 300 mM NaCl, 0.1% (v/v) Triton X-100, 1 mM DTT, 1 mM benzamidine, 1 mM PMSF, lysozyme, DNAse, RNAse and protease inhibitor (Sigmafast)) and lysed (Branson Sonifier 250). Debris was removed by centrifugation and the supernatant applied to a Ni-NTA gravity flow column (Qiagen). After washing (20 mM Tris pH 8, 20 mM imidazole, 300 mM NaCl), proteins were eluted with 500 mM imidazole. Protein containing fractions were dialyzed overnight (20 mM Tris pH 7 (pH 8 for IDR^Δ97^), 300 mM NaCl, 10% v/v glycerol, 1 mM DTT, 0.5 mM PMSF) in the presence of Ulp-1 protease in a molar ratio of 20:1 to yield the native TRPV4 N-terminal constructs. After dialysis, cleaved proteins were separated by a reverse Ni-NTA affinity chromatography step and subsequently purified via a HiLoad prep grade 16/60 Superdex200 or 16/60 Superdex75 column (GE Healthcare) equilibrated with 20 mM Tris pH 7, 300 mM NaCl, 1 mM DTT. Pure sample fractions were flash-frozen in liquid nitrogen and stored at −20 °C until further use.

Purified IDR^Δ97^ was extensively dialyzed against double distilled water, lyophilized, and stored in solid form at −20 °C. Peptides could be dissolved in desired amounts of buffer to concentrations up to 10 mM.

### Analytical size-exclusion chromatography

Analytical SEC experiments were carried out at 4 °C using an NGC Quest (BioRad) chromatography system. 250 μL protein at a concentration of 2-3 mg/mL was injected on a Superdex200 10/300 increase column (GE Healthcare) equilibrated with 20 mM Tris pH 7, 300 mM NaCl, 1 mM DTT via a 1 mL loop. Protein was detected by absorbance measurement at wavelengths of 230 and 280 nm.

For Stokes radius (*R_S_*) determination, SEC columns were calibrated with a protein standard kit (GE Healthcare) containing ferritin (*MW* = 440 kDa, *R_S_* = 61.0 nm), alcohol dehydrogenase (150 kDa, 45.0 nm), conalbumin (75 kDa, 36.4 nm), ovalbumin (43 kDa, 30.5 nm), carbonic anhydrase (29 kDa, 23.0 nm), ribonuclease A (13.7 kDa, 16.4 nm), and aprotinin (6.5 kDa, 13.5 nm) whose Stokes radii were obtained from La Verde et al.^56^ The SEC elution volume, *V_e_* (in mL), of the protein standards was plotted versus the log(*R_S_*), with *R_S_* in nm, and fitted with a linear regression (Equation **1**):

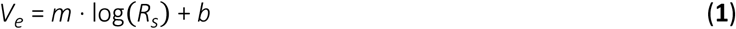

where *m* is the slope of the linear regression and *b* the *y*-axis section. Equation **1** was then used to calculate the Stokes radii of the TRPV4 constructs from their respective SEC elution volumes.

### Size exclusion chromatography multi-angle light scattering (SEC-MALS)

Multi-angle light scattering coupled with size-exclusion chromatography (SEC-MALS) of the *G. gallus* TRPV4 NTD, ARD, and IDR was performed with a GE Superdex200 Increase 10/300 column run at 0.5 mL/min on a Jasco HPLC unit (Jasco Labor und Datentechnik) connected to a light scattering detector measuring at three angles (miniDAWN TREOS, Wyatt Technology). The column was equilibrated for at least 16 hrs with 20 mM Tris pH 7, 300 mM NaCl, 1 mM DTT (filtered through 0.1 μm pore size VVLP filters (Millipore)) before 200 μL of protein samples at a concentration of 2 mg/mL were loaded. The ASTRA software package (Wyatt Technology) was used for data analysis, assuming a Zimm model^57^. The molecular weight, *M_W_*, can be determined from the reduced Rayleigh ratio extrapolated to zero, *R*(0), which is the light intensity scattered from the analyte relative to the intensity of the incident beam (Equation **2**):

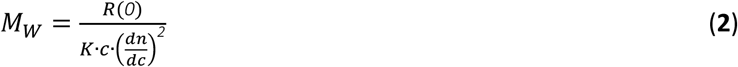

Here, *c* is the concentration of the analyte and (*dn*/*dc*) is the refractive index increment, which was set to 0.185 mL/g, a standard value for proteins^58^. *K* is an optical constant depending on wavelength and the solvent refractive index. The protein extinction coefficients at 280 nm were calculated from the respective amino acid sequences using the ProtParam tool^59^.

### Circular dichroism (CD) spectroscopy

CD measurements were carried out on a Jasco-815 CD spectrometer (JascoTM) with 1 mm quartz cuvettes (Hellma Macro Cell). Proteins were used at concentrations in the range of 0.03-0.05 mg/mL in 5 mM Tris pH7, 10 mM NaCl. Spectra were recorded at 20 °C between 190 and 260 nm with 1 nm scanning intervals, 5 nm bandwidth and 50 nm/min scanning speed. All spectra were obtained from the automatic averaging of three measurements with automatic baseline correction. The measured ellipticity *θ* in degrees (deg) was converted to the mean residue ellipticity (MRE) via equation **3**^60^.

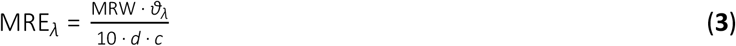

Here, MRE_λ_ is the mean residue ellipticity, and *θ_λ_* is the measured ellipticity at wavelength *λ*, *d* is the pathlength (in cm), and *c* is the protein concentration (g/mL). MRW is the mean residue weight, MRW = *M_W_* ・ (*N*-1)-1, where *M_W_* is the molecular weight of the protein (in Da), and *N* is the number of residues. For titration experiments, TRPV4 N-terminal peptides were used in a concentration of 30 μM in double distilled water in the presence of TFE (2,2,2-trifluoroethanol, 0-90% (v/v)), SDS (0.5, 1.0, 2.5, 5.0 and 8.0 mM) and liposomes (0.5 and 1.0 mM). Liposomes were prepared from POPG and POPC at a molar ratio of 1:1 as described below.

### Small angle X-ray scattering (SAXS)

SAXS experiments were carried out at the EMBL-P12 bioSAXS beam line, DESY^61^. SEC-SAXS data collection^62^, *I*(*q*) vs *q*, where *q* = 4*π*sin*θ*/*λ*; 2*θ* is the scattering angle and *λ* the X-ray wavelength (0.124 nm; 10 keV) was performed at 20 °C using S75 (IDR constructs) and S200 Increase 5/150 (NTD and ARD constructs) analytical SEC columns (GE Healthcare) equilibrated in the appropriate buffers (see Tables S1 and S2) at flow rates of 0.3 mL/min. Automated sample injection and data collection were controlled using the *BECQUEREL* beam line control software^63^. The SAXS intensities were measured as a continuous series of 0.25 s individual X-ray exposures, from the continuously-flowing column eluent, using a Pilatus 6M 2D-area detector for a total of one column volume (ca. 600-3000 frames in total). The 2D-to-1D data reduction, i.e., radial averaging of the data to produce 1D *I*(*q*) vs *q* profiles, were performed using the SASFLOW pipeline incorporating RADAVER from the ATSAS 2.8 suite of software tools^64^. The individual frames obtained for each SEC-SAXS run were processed using CHROMIXS^65^. Briefly, individual SAXS data frames were selected across the respective sample SEC-elution peaks and an appropriate region of the elution profile, corresponding to SAXS data measured from the solute-free buffer, were identified, averaged and then subtracted to generate individual background-subtracted sample data frames. These data frames underwent further CHROMIXS analysis, including the assessment of the radius of gyration (*R_g_*) of each individual sample frame, scaling of frames with equivalent *R_g_*, and subsequent averaging to produce the final 1D-reduced and background-corrected scattering profiles. Only those scaled individual SAXS data frames with a consistent *R_g_* through the SEC-elution peak that were also evaluated as statistically similar through the measured *q*-range were used to generate the final SAXS profiles. Corresponding UV traces were not measured; the column eluate was flowed directly to the P12 sample exposure unit after the small column, forging UV absorption measurements, to minimize unwanted band-broadening of the sample. All SAXS data-data comparisons and data-model fits were assessed using the reduced *c*^2^ test and the Correlation Map, or CORMAP, *p*-value^66^. Fits within the *c*^2^ range of 0.9–1.1 or having a CORMAP *p*-values higher than the significance threshold cutoff of a = 0.01 are considered excellent, i.e., no systematic differences are present between the data-data or data-model fits at the significance threshold.

Primary SAXS data analysis was performed using PRIMUS as well as additional software modules from ATSAS 3.0.1^67^. The Guinier approximation^68^ (ln(*I*(*q*)) vs. *q*^2^ for *qR_g_* < 1.3) and the real-space pair distance distribution function, or *p*(*r*) profile (calculated from the indirect inverse Fourier transformation of the data, thus also yielding estimates of the maximum particle dimension, *D_max_*, Porod volume, *V_p_*, shape classification, and concentration-independent molecular weight^69–71^ were used to estimate the *R_g_* and the forward scattering at zero angle, *I*(0). Dimensionless Kratky plot representations of the SAXS data (*qR_g_*^2^(*I*(*q*)/*I*(0)) vs. *qR_g_*) followed an approach previously described^72^. All collected SAXS data are reported in Tables S1 and S2.

### DAMMIN modeling

The shape reconstruction of the ARD was performed using DAMMIN^36^ where nine individual dummy-atom models that fit the SAXS data underwent spatial alignment with DAMSEL and DAMSUP, followed by volume and bead occupancy correction with DAMAVER/DAMFILT/DAMSTART^73^ to generate a final overall shape of the protein.

### Rigid body and ensemble modeling

Subsequent rigid-body normal mode analysis of the ARD was performed using the program SREFLEX^74^ using the X-ray crystal structure (PDB: 3W9G) as a template. CRYSOL was used to assess data-model fits^75^. The ensemble analysis of IDR, NTD, the systematic NTD IDR-deletions and/or respective IDR/NTD-PIP_2_-binding site mutants was performed using Ensemble Optimization Method, EOM^34,35^. Briefly, 10000 protein structures were generated for each of the respective protein constructs, where the IDR section(s) were modelled as random chains (self-avoiding walks with the confines of Ramachandran-constraints). The scattering profiles were calculated for each model within the initially generated 10000 member ensembles. The selection of sub-ensembles describing the SAXS data, and the assessment of the *R_g_* distribution of the refined ensemble pools, was performed using a genetic algorithm based on fitting the SAXS data with a combinatorial volume-fraction weighted sum contribution of individual model scattering profiles drawn from the initial pool of structures.

### Hydrogen/deuterium exchange mass spectrometry (HDX-MS)

HDX-MS was conducted on three independent preparations of *G. gallus* TRPV4 IDR, ARD or NTD protein each, and for each of those three technical replicates (individual HDX reactions) per deuteration timepoint were measured. Preparation of samples for HDX-MS was aided by a two-arm robotic autosampler (LEAP Technologies). HDX reactions were initiated by 10-fold dilution of the proteins (25 μM) in buffer (20 mM Tris pH 7, 300 mM NaCl) prepared in D2O and incubated for 10, 30, 100, 1,000 or 10,000 s at 25 °C. The exchange was stopped by mixing with an equal volume of pre-dispensed quench buffer (400 mM KH_2_PO_4_/H_3_PO_4_, 2 M guanidine-HCl; pH 2.2) kept at 1 °C, and 100 μl of the resulting mixture injected into an ACQUITY UPLC M-Class System with HDX Technology^76^. Non-deuterated samples were generated by a similar procedure through 10-fold dilution in buffer prepared with H_2_O. The injected HDX samples were washed out of the injection loop (50 μL) with water + 0.1% (v/v) formic acid at a flow rate of 100 μL/min and guided over a column containing immobilized porcine pepsin kept at 12 °C. The resulting peptic peptides were collected on a trap column (2 mm x 2 cm), that was filled with POROS 20 R2 material (Thermo Scientific) and kept at 0.5 °C. After three minutes, the trap column was placed in line with an ACQUITY UPLC BEH C18 1.7 μm 1.0 x 100 mm column (Waters) and the peptides eluted with a gradient of water + 0.1% (v/v) formic acid (eluent A) and acetonitrile + 0.1% (v/v) formic acid (eluent B) at 60 μL/min flow rate as follows: 0-7 min/95-65% A, 7-8 min/65-15% A, 8-10 min/15% A. Eluting peptides were guided to a Synapt G2-Si mass spectrometer (Waters) and ionized by electrospray ionization (capillary temperature and spray voltage of 250 °C and 3.0 kV, respectively). Mass spectra were acquired over a range of 50 to 2,000 *m/z* in enhanced high definition MS (HDMS^E^)^77,78^ or high definition MS (HDMS) mode for non-deuterated and deuterated samples, respectively. Lock mass correction was conducted with [Glu1]-Fibrinopeptide B standard (Waters). During separation of the peptides on the ACQUITY UPLC BEH C18 column, the pepsin column was washed three times by injecting 80 μL of 0.5 M guanidine hydrochloride in 4% (v/v) acetonitrile. Blank runs (injection of double-distilled water instead of the sample) were performed between each sample. All measurements were carried out in triplicates. Peptides were identified and evaluated for their deuterium incorporation with the software ProteinLynx Global SERVER 3.0.1 (PLGS) and DynamX 3.0 (both Waters). Peptides were identified with PLGS from the non-deuterated samples acquired with HDMS^E^ employing low energy, elevated energy and intensity thresholds of 300, 100 and 1,000 counts, respectively and matched using a database containing the amino acid sequences of IDR, ARD, NTD, porcine pepsin and their reversed sequences with search parameters as follows: Peptide tolerance = automatic; fragment tolerance = automatic; min fragment ion matches per peptide = 1; min fragment ion matches per protein = 7; min peptide matches per protein = 3; maximum hits to return = 20; maximum protein mass = 250,000; primary digest reagent = non-specific; missed cleavages = 0; false discovery rate = 100. For quantification of deuterium incorporation with DynamX, peptides had to fulfil the following criteria: Identification in at least 2 of the 3 non-deuterated samples; the minimum intensity of 10,000 counts; maximum length of 30 amino acids; minimum number of products of two; maximum mass error of 25 ppm; retention time tolerance of 0.5 minutes. All spectra were manually inspected and omitted if necessary, e.g. in case of low signal-to-noise ratio or the presence of overlapping peptides disallowing the correct assignment of the isotopic clusters.

Residue-specific deuterium uptake from peptides identified in the HDX-MS experiments was calculated with the software DynamX 3.0 (Waters). In the case that any residue is covered by a single peptide, the residue-specific deuterium uptake is equal to that of the whole peptide. In the case of overlapping peptides for any given residue, the residue-specific deuterium uptake is determined by the shortest peptide covering that residue. Where multiple peptides are of the shortest length, the peptide with the residue closest to the peptide C-terminus is utilized. Assignment of residues being intrinsically disordered was based on two criteria, i.e., a residue-specific deuterium uptake of >50% after 10 s of HDX and no further increment in HDX >5% in between consecutive HDX times. Raw data of deuterium uptake by the identified peptides and residue-specific HDX are provided in Supplemental Dataset 1.

### Crosslinking mass spectrometry (XL-MS)

For structural analysis, 100 μg of purified protein were crosslinked by addition of DSS-H12/D12 (Creative Molecules) at a ratio of 1.5 nmol / 1 μg protein and gentle shaking for 2 h at 4 °C. The reaction was performed at a protein concentration of 1 mg/mL in 20 mM HEPES pH 7, 300 mM NaCl. After quenching by addition of ammonium bicarbonate (AB) to a final concentration of 50 mM, samples were dried in a vacuum centrifuge. Then, proteins were denatured by resuspension in 8M urea, reduced with 2.5 mM Tris(2-carboxylethyl)-phosphine (TCEP) at 37 °C for 30 min and alkylated with 5 mM iodoacetamide at room temperature in the dark for 30 min. After dilution to 1 M urea using 50 mM AB, 2 μg trypsin (protein:enzyme ratio 50:1; Promega) were added and proteins were digested at 37 °C for 18 h. The resulting peptides were desalted by C18 Sep-Pak cartridges (Waters), then crosslinked peptides were enriched by size exclusion chromatography (Superdex 20 Increase 3.2/300, cytiva) prior to liquid chromatography (LC)-MS/MS analysis on an Orbitrap Fusion Tribrid mass spectrometer (Thermo Scientific). MS measurement was performed in data-dependent mode with a cycle time of 3 s. The full scan was acquired in the Orbitrap at a resolution of 120,000, a scan range of 400-1500 m/z, AGC Target 2.0e5 and an injection time of 50 ms. Monoisotopic precursor selection and dynamic exclusion for 30 s were enabled. Precursor ions with charge states of 3-8 and minimum intensity of 5e3 were selected for fragmentation by CID using 35% activation energy. MS2 was done in the Ion Trap in rapid scan range mode, AGC target 1.0e4 and a dynamic injection time. All experiments were performed in biological triplicates and samples were measured in technical duplicates. Crosslink analysis was done with the *xQuest/xProphet* pipeline^79^ in ion-tag mode with a precursor mass tolerance of 10 ppm. For matching of fragment ions, tolerances of 0.2 Da for common ions and 0.3 Da for crosslink ions were applied. Crosslinks were only considered for further analyses if they were identified in at least 2 of 3 biological replicates with deltaS < 0.95 and at least one Id score ≥ 25.

### Lipid preparation

Liposomes were prepared from 1-palmitoyl-2-oleoyl-*sn*-glycero-3-phosphocholine (POPC) and 1-palmitoyl-2-oleoyl-*sn*-glycero-3-phosphoglycerol (POPG) mixed in a 1:1 (n/n) ratio in chloroform. The organic solvent was removed via nitrogen flux and under vacuum via desiccation overnight. The lipid cake was suspended in 1 mL buffer (10 mM Tris pH 7, 100 mM NaCl or 300 mM NaCl) and incubated for 20 min at 37°C and briefly spun down before being subjected to five freeze and thaw cycles. The resulting large unilamellar vesicles (LUVs) were incubated for 20 min at 21 °C under mild shaking. To obtain a homogeneous solution of small unilamellar vesicles (SUVs), the mixture was extruded 15 times through a 100 nm membrane using the Mini Extruder (Avanti Polar Lipids). This yielded a liposome stock solution of 100 nm liposomes with 4.0 mM lipid in 10 mM Tris pH 7, 100 mM NaCl that was used immediately for measurements. POPC-only liposomes were prepared similarly. 1,2-dioctanoyl-*sn*-glycero-3-phospho-(1’-*myo*-inositol-4’,5’-bisphosphate) (diC8-PI(4,5)P_2_) was purchased as a powder and directly dissolved in the appropriate amount of buffer.

### Liposome sedimentation assay

Liposomes were prepared as described above. Proteins and liposomes in 20 mM Tris pH 7, 100 mM NaCl were mixed to a final concentration of 2.5 μM protein and 2 mg/mL lipid. After incubation at 4°C for 1 hr under mild shaking, an SDS-PAGE sample of the input was taken. The mixture was then centrifuged at 70,000 g for 1 hr at 4 °C. SDS-PAGE samples (15 μL) were taken from both the supernatant and the pellet resuspended in assay buffer. Control samples without liposomes were run in parallel to verify the protein stability under the experimental conditions. The protein distribution between the pellet and supernatant fractions was determined by running an SDS-PAGE and densitometrically analyzing the bands using imageJ^53^. Sedimentation assays were carried out three times for each protein liposome mixture and protein only control sample. Error bars were calculated as the standard deviation from the mean value of three replicates.

### Nuclear magnetic resonance (NMR) spectroscopy

Backbone assignments of native ^13^C, ^15^N-labeled *G. gallus* IDR have been reported by us previously^5^. For complete backbone assignments of IDR^AAWAA^ and IDR^Patch^, ^15^N, ^13^C-labeled proteins were prepared. Backbone and side chain chemical shift resonances were assigned with a set of band-selective excitation short-transient (BEST) transverse relaxation-optimized spectroscopy (TROSY)-based assignment experiments: HNCO, HN(CA)CO, HNCA, HN(CO)CA, HNCACB. Additional side chain chemical shift information was obtained from H(CCCO)NH and (H)CC(CO)NH experiments.

For titrations with lipids, or comparison of chemical shifts between constructs, ^15^N-labeled IDR and NTD variants were prepared. All NMR spectra were recorded at 10 °C on 600 MHz to 950 MHz Bruker AvanceIII HD NMR spectrometer systems equipped with cryogenic triple resonance probes. For peptide and lipid titration experiments, a standard [^1^H,^15^N]-BEST-TROSY pulse sequence implemented in the Bruker Topspin pulse program library was used. Solutions with 100 μM of ^15^N-labeled IDR constructs in 10 mM Tris-HCl pH 7, 100 mM (or 300 mM) NaCl, 1 mM DTT, 10% (v/v) D2O were titrated with lipid from a concentrated stock solution. The chemical shifts were determined using TopSpin 3.6 (Bruker) The ^1^H and ^15^N weighted chemical shift differences observed in ^1^H, ^15^N-HSQC spectra were calculated according to equation **4**^80^:

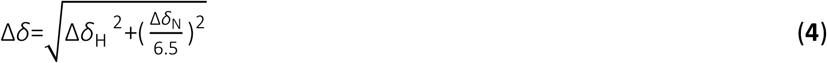

Here, Δ*δ*_H_ is the ^1^H chemical shift difference, Δ*δ*_N_ is the ^15^N chemical shift difference, and Δ*δ* is the ^1^H and ^15^N weighted chemical shift difference in ppm.

For NMR titrations of ^15^N-labeled TRPV4 IDR with liposomes (SUVs) where line broadening instead of peak shifts was observed, the interaction of the reporter with liposomes was quantified using the peak signal loss in response to liposome titration. The signal loss at a lipid concentration *ci* was calculated as the relative peak signal decrease rel. Δ*I* according to equation **5**.

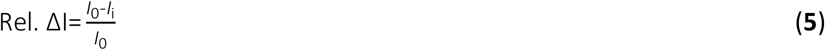

Here, *I_0_* is the peak integral in the absence of SUVs, and *I_i_* is the peak integral in the presence of a lipid concentration *c_i_*.

^31^P{^1^H} NMR spectra were recorded at 25 °C on a 600 MHz Bruker AvanceIII HD NMR spectrometer. DiC_8_-PI(_4,5_)P_2_ was used at 500 μM in 10 mM Tris pH 7, 100 mM NaCl, 10% (v/v) D_2_O and titrated with protein from a concentrated stock solution.

### Tryptophan fluorescence spectroscopy

All tryptophan fluorescence measurements were carried out in 10 mM Tris pH 7, 100 mM NaCl buffer on a Fluro Max-4 fluorimeter with an excitation wavelength of 280 nm and a detection range between 300 nm to 550 nm. The fluorescence wavelength was determined as the intensity-weighted fluorescence wavelength between 320-380 nm (hereafter referred to as the average fluorescence wavelength) according to equation **6**:

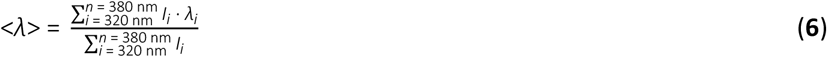

Here, <*λ*> is the average fluorescence wavelength and *I_i_* is the fluorescence intensity at wavelength *λ_i_*.

To monitor liposome binding by tryptophan fluorescence, fluorescence emission spectra were recorded in the presence of increasing lipid concentrations. Lipids were prepared as SUVs with 100 nm diameters as described above. The protein concentrations were kept constant at 5 μM. The protein-liposome mixtures were incubated for 10 min prior to recording the emission spectra. For each sample, at least three technical replicates were measured. The tryptophan fluorescence wavelength at each lipid concentration was quantified by determining the average fluorescence wavelength using equation (**6**). The dissociation constants, *K_d_*, of the protein liposome complexes were determined by plotting the changes in <*λ*> against the lipid concentration *c* and fitting the data with a Langmuir binding isotherm (equation (**7**)):

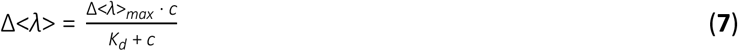

Here, Δ<*λ*> describes the wavelength shift between the spectrum in the absence of lipids and a given titration step at a lipid concentration *c*. Δ<*λ*>_*max*_ indicates the maximum wavelength shift in the saturation regime of the binding curve. Under the assumption that proteins cannot diffuse through liposome membranes and therefore can only bind to the outer leaflet, the lipid concentration *c* was set to half of the titrated lipid concentration.

### Calcium imaging

MN-1 cells were transfected with GFP-tagged TRPV4 plasmids using Lipofectamine LTX with Plus Reagent. Calcium imaging was performed 24 h after transfection on a Zeiss Axio Observer.Z1 inverted microscope equipped with a Lambda DG-4 (Sutter Instrument Company, Novato, CA) wavelength switcher. Cells were bath-loaded with Fura-2 AM (8 μM, Life Technologies) for 45-60 min at 37°C in calcium-imaging buffer (150 mM NaCl, 5 mM KCl, 1 mM MgCl_2_, 2 mM CaCl_2_, 10 mM glucose, 10 mM HEPES, pH 7.4). For hypotonic saline treatment, one volume of NaCl-free calcium-imaging buffer was added to one volume of standard calcium-imaging buffer for a final NaCl concentration of 70 mM. For GSK101 treatment, GSK101 was added directly to the calcium imaging buffer to achieve 50 nM final concentration. Cells were imaged every 10 s for 20 s prior to stimulation with hypotonic saline or GSK101, and then imaged every 10 s for an additional 2 min. Calcium levels at each time point were computed by determining the ratio of Fura-2 AM emission at 340 nM divided by the emission at 380 nM. Data were expressed as raw Fura ratio minus background Fura ratio.

### Molecular dynamics (MD) simulations

All simulations were performed using Gromacs 2020.3^81^ and the MARTINI2.2 forcefield^82,83^ with rescaled protein-protein interactions to better represent the disordered nature of the IDR^84^. The scaling factor was set to *α* = 0.87, which best describes the measured R*g* distribution of the native IDR (Fig. S14). Protein-membrane interactions were not rescaled. All production simulations were performed with a 20 fs integration timestep. For equilibration simulations of systems with only protein, water, and NaCl ions present, a 40 fs timestep was used. A temperature of 37 °C was maintained with thermostats acting on protein, membrane, and solvent (water and ions) individually. The Berendsen thermostat^85^ was used for equilibration simulations and the v-rescale thermostat^86^ for production simulations, in both cases with characteristic times of 1 ps. A pressure of 1 bar was established with a semi-isotropic barostat (with coupled x and y dimensions) in simulations with a membrane present and with an isotropic barostat otherwise. We employed the Berendsen barostat^85^ for equilibration simulations and switched to the Parrinello-Rahman barostat^87^ for production simulations, always using a 20 ps time constant and a compressibility factor of 3 × 10^−4^ bar^−1^. Bond constraints were maintained using the LINCS algorithm.^88^ To alleviate unequal heating of different lipid types, we increased the default LINCS order to 8.^89^ Electrostatic interactions and van-der-Waals interactions were cut-off at 1.1 nm. All simulations were performed with an increased cut-off distance of 1.418 nm for the short-range neighbour list. Replicates were started from the same equilibrated structures, but their initial velocities were independently drawn from the Maxwell-Boltzmann distribution for each replicate simulation.

To set up the simulation systems, native IDR, IDR^AAWAA^ and IDR^Patch^ were modelled as disordered coils with atomistic resolution using the VMD molefacture plugin^90^, converted to coarse-grained topologies using the martinize.py script (version 2.6), and placed in a 30 x 30 x 30 nm^3^ box with solvent and 150 mM NaCl. The systems were then energy minimized using a steepest descent algorithm for 3000 steps.

Subsequently, the systems were equilibrated for 20 ns with downscaled protein-protein interactions (*α* = 0.3; only used during this step) to generate a relatively open initial IDR structure. The protein structure was then extracted and placed in a random position in the water phase of a 20 x 20 x 20 nm^3^ box that contained a preequilibrated patch of a membrane modelled after the inner leaflet of the plasma membrane (see Table S3 for membrane composition). Steepest descent energy minimization for 1000 steps was followed by MD equilibration for 100 ns. To probe the effect of IDR positioning in the full TRPV4 assembly, we set up simulations in which the backbone bead of V134 (C-terminus of the IDR) was harmonically restrained to a height of |*z*(V134)-*z*(membrane)|=5, 6, 7, 8 or 9 nm over the midplane of the membrane with a force constant of 1000 kJ mol^−1^ nm^−2^. Here, *z*(membrane) is the center of mass of the membrane. Five sets of simulations (comprising four replicates each) were carried out for each of the three constructs using the Gromacs pull code^81^. To emulate the NMR experiments, we also performed four replicate MD simulations for each IDR construct without distance restraint. Each replicate was simulated for approximately 38 μs. After 10 μs full membrane binding was obtained in all systems and only the following approximately 28 μs of each replicate were considered for analysis. In one simulation with IDR^Patch^ restrained at 9 nm, the IDR reached over the periodic boundary to the other face of the membrane. This simulation was hence not included in any analysis. VMD^90^ was used for visual analysis and rendering. All analyses were carried out with python scripts.

## Data availability

The NMR backbone assignment of the *G. gallus* TRPV4 N-terminal intrinsically disordered region has been deposited in the BioMagResBank (www.bmrb.io) under the accession number 51172. The SAXS data have been deposited in the SASBDB under the accession numbers SASDQE8 (ARD), SASDQF8 (NTD), SASDQG8 (NTD^AAWAA^), SASDQH8 (NTD^ΔN54^), SASDQJ8 (NTD^ΔN97^), SASDQK8 (NTD^ΔN104^), SASDQL8 (NTD^ΔN120^), SASDQM8 (IDR), SASDQN8 (IDR^AAWAA^). A summary of the conditions used for HDX-MS analyses and a full list of the peptides obtained for different TRPV4 protein constructs is available in Supplemental Dataset 1. The XL-MS data have been deposited to the ProteomeXchange Consortium via the PRIDE partner repository^91^ with the project accession number PXD038153, a summary of the peptides identified by XL-MS is available in Supplemental Dataset 2.

## Acknowledgements

We acknowledge the generous support by the beamline staff scientists at EMBL/P12. We thank Sabine Häfner for technical support, Rupert Abele and Andreas Schlundt for help with SEC-MALS and SAXS measurements, Marta Bogacz and Lisa Pietrek for fruitful discussions, and Ainara Claveras Cabezudo for help in TRPV4 modeling. This project was supported by the Centre for Biomolecular Magnetic Resonance (BMRZ), Goethe University Frankfurt, funded by the state of Hesse. Access to beamline P12, EMBL (DESY), Hamburg was made available via iNEXT-ERIC, BAG proposal #SAXS-1106 “Conformational dynamics and equilibria in regulatory multi-domain proteins, RNAs and their complexes” (to UAH). BG acknowledges a PhD fellowship of the Max Planck Graduate Center (MPGC). This project received funding from the NIH (NINDS K08 NS102509 to BAM, R35 NS122306 to CJS), the Muscular Dystrophy Association (project 629305 to CJS), the core facility for Interactions, Dynamics and Macromolecular Assembly (WS, project 324652314 to Gert Bange, Marburg), the Max Planck Society (SLS and GH), and the Deutsche Forschungsgemeinschaft (DFG, German Research Foundation) through grant STE 2517/5-1 (to FS), the collaborative research center 1507 “Membrane-associated Protein Assemblies, Machineries, and Supercomplexes“ – Project ID 450648163 (to UAH and GH) and the Cluster of Excellence “Balance of the Microverse” EXC 2051 – Project-ID 390713860 (to UAH). UAH acknowledges an instrumentation grant for a high-field NMR spectrometer by the REACT-EU EFRE Thuringia (Recovery assistance for cohesion and the territories of Europe, European Fonds for Regional Development, Thuringia) initiative of the European Union.

## Author contributions

Conceptualization: B.G., U.A.H., analysis: B.G., C.W. B.A.M., S.L.S., J.J., C.M.J., W.S., G.H., U.A.H.; investigation: B.G., C.W. B.A.M., S.L.S., J.J., F.T., S-A.M., J.N., J.K.D., C.M.J., W.S.; writing – original draft: B.G., U.A.H., writing – review and editing: B.G., C.W., S.L.S., G.H., U.A.H.; visualization: B.G., C.W., S.L.S., U.A.H.; supervision: B.A.M., F.S., C.J.S., G.H., U.A.H.; funding acquisition: B.A.M., W.S., F.S., C.J.S., G.H., U.A.H. All authors read and approved the final version of the manuscript.

## Additional information

Supplemental Dataset 1. Summary of HDX-MS analyses and full list of the peptides obtained for different TRPV4 protein constructs.

Supplemental Dataset 2. Summary of XL-MS analyses.

Movie S1 – Ensemble of TRPV4 IDR structures from MARTINI Simulations on the membrane_side view.

Movie S2 - Ensemble of TRPV4 IDR structures from MARTINI Simulations on the membrane_bottom view.

Correspondence and requests for materials should be addressed to U.A.H.

