## Supporting Information for "Crosstalk between regulatory elements in the disordered TRPV4 N-terminus modulates lipid-dependent channel activity"

**Supporting Information**
**for**

### Crosstalk between regulatory elements in the disordered TRPV4 N-terminus modulates lipid-dependent channel activity

Benedikt Goretzki[1,2], Christoph Wiedemann[1], Brett A. McCray[3], Stefan L. Schäfer[4], Jasmin Jansen[5,6], Frederike Tebbe[1], Sarah-Ana Mitrovic[7], Julia Nöth[7], Jack K. Donohue[3], Cy M. Jeffries[8], Wieland Steinchen[9], Florian Stengel[5,6], Charlotte J. Sumner[3,10], Gerhard Hummer[4,11], Ute A. Hellmich[1,2,*]

[1]Friedrich Schiller University Jena, Faculty of Chemistry and Earth Sciences, Institute of Organic Chemistry and Macromolecular Chemistry, Humboldtstraße 10, 07743 Jena, Germany

[2]Centre for Biomolecular Magnetic Resonance (BMRZ), Goethe University, Max von Laue Str. 9, 60438 Frankfurt, Germany

[3]Department of Neurology, Johns Hopkins University School of Medicine, Baltimore, MD, USA

[4] Department of Theoretical Biophysics, Max Planck Institute of Biophysics, Max-von-Laue Str. 3, 60438 Frankfurt am Main, Germany

[5]Department of Biology, University of Konstanz, Universitätsstraße 10, 78457 Konstanz, Germany

[6]Konstanz Research School Chemical Biology, University of Konstanz, Universitätsstraße 10, 78457 Konstanz, Germany

[7]Department of Chemistry, Section Biochemistry, Johannes Gutenberg-University Mainz, Joachim-Becher-Weg 30, 55128 Mainz, Germany

[8]European Molecular Biology Laboratory, EMBL Hamburg Unit, Deutsches Elektronen-Synchrotron, Notkestraße 85, 22607 Hamburg, Germany

[9]Center for Synthetic Microbiology (SYNMIKRO) & Department of Chemistry, Philipps-University Marburg, Karl-von-Frisch-Str. 14, 35043 Marburg, Germany

[10]Department of Neuroscience, Johns Hopkins University School of Medicine, Baltimore, MD, USA

[11]Institute of Biophysics, Goethe University Frankfurt, Max-von-Laue Str. 1, 60438 Frankfurt am Main, Germany

**Table S1 – SAXS data reporting table for native TRPV4 NTD, ARD and IDR as well as $PIP_2$-binding site mutants.**

| | | | | | |
|---|---|---|---|---|---|
| **Sample details** | | | | | |
| SAMPLE | NTD | $NTD^{AAWAA}$ | ARD | IDR | $IDR^{AAWAA}$ |
| SASBDB Accession Codes | | | | | |
| Organism | | | *Gallus gallus* | | |
| NCBI protein accession ID | | | 395427 | | |
| (amino acid range) | 2-382 | 2-382 | 135-382 | 2-134 | 2-134 |
| SEC-SAXS buffer | | | 20 mM Tris pH 7.0, 10 mM DTT | | |
| NaCl concentration | | 300 mM | | 100 mM | |
| Sample injection volume | 40 µl | 40 µl | 40 µl | 40 µl | 40 µl |
| Sample injection conc. | 10.0 mg/ml | 8.2 mg/ml | 10.7 mg/ml | 10.1 mg/ml | 12.9 mg/ml |
| SEC column | | S200 Increase 5/150 | | S75 Increase 5/150 | |
| SEC flow rate | | | 0.3 ml/min | | |
| SEC temperature | | | 20 °C | | |
| **Instrument details** | | | | | |
| Instrument | | | EMBL P12 bioSAXS beam line, DESY, Hamburg | | |
| Exposure time/# frames | | | 0.25 s (2400) | | |
| X-ray wavelength/energy | | | 0.124 nm (9996.5 eV) | | |
| Sample-to-detector distance | | | 3 m | | |
| Scattering intensity scale | | | Arbitrary unit, a.u. | | |
| SEC-SAXS primary data processing | | | CHROMIXS (ATSAS 3.0.1) | | |
| # frames used for averaging | 104 | 95 | 79 | 59 | 60 |
| Working s-range ($nm^{-1}$) | 0.07-3.70 | 0.06-4.20 | 0.09-4.98 | 0.08-5.00 | 0.10-5.60 |
| **Guinier analysis:** | | | | | |
| Primary data analysis software | | | PRIMUS (ATSAS 3.0.1) | | |
| Guinier I(0) ($\sigma$) | 1721(8) | 5002(13) | 3600(5) | 1997(9) | 4629(10) |
| $R_g$ (Guinier, nm) ($\sigma$) | 3.68(0.03) | 4.02(0.01) | 2.33(0.01) | 3.20(0.02) | 3.26(0.02) |
| $sR_g$ range | 0.26-1.20 | 0.40-1.01 | 0.21-1.30 | 0.25-1.30 | 0.35-1.29 |
| **p(r) analysis:** | | | | | |
| Method | | | GNOM 5 | | |
| I(0), POR ($\sigma$) | 1754(11) | 5147(17) | 3644(9) | 2009(11) | 4714(12) |
| $R_g$ (POR, nm) ($\sigma$) | 4.06(0.07) | 4.53(0.04) | 2.49(0.02) | 3.44(0.05) | 3.50(0.02) |
| $D_{max}$ (nm) | 19.0 | 19.5 | 11.5 | 14.5 | 14.5 |
| Quality of fit, CorMap P / $\chi^2$ | 0.62/1.03 | 0.69/0.99 | 0.50/1.07 | 0.90/0.98 | 0.26/1.07 |
| Porod volume ($nm^3$) | 59 | 78 | 41 | 58 | 37 |
| Shape classification | flexible | flexible | compact | random chain | random chain |
| **Molecular Weight analysis:** | | | | | |
| MW, calculated from amino acid sequence, kDa | 42.5 | 42.2 | 28.0 | 14.5 | 14.2 |
| MW from SAXS data, kDa | 38-42 | 48-56 | 26-29 | 24-29 | 22-25 |
| **Ab initio modelling:** | | | | | |
| Method | | | DAMMIN | | |
| Symmetry | | | P1 | | |
| #models used for averaging | | | 9 | | |
| Normalized Spatial Discrepancy, NSD | | | 0.58 | | |
| Resolution estimate, nm | | | 24 | | |
| Quality-of-fit, CorMap P / $\chi^2$ | | | 0.45/1.06 | | |
| **Rigid body/Normal mode modelling:** | | | | | |
| Method | | | SREFLEX | | |
| Symmetry | | | P1 | | |
| Template | | | PDB ID: 3W9G | | |
| Initial template fit, CorMap P / $\chi^2$ | | | 1.44e-43/3.33 | | |
| Final model fit, CorMap P / $\chi^2$ | | | 4.05e-07/1.21 | | |
| **Ensemble modelling:** | | | | | |
| Method | EOM | EOM | | EOM | EOM |
| Symmetry | P1 | P1 | | P1 | P1 |
| Template | PDB ID: 3W9G | PDB ID: 3W9G | | - | - |
| Final model fit, CorMap P / $\chi^2$ | 0.72/0.99 | 0.52/0.99 | | 0.97/0.98 | 0.19/1.3 |

**Table S2 – SAXS data reporting table for TRPV4 NTD deletion constructs.**

| Sample details | | | | |
|---|---|---|---|---|
| SAMPLE | $NTD^{\Delta N120}$ | $NTD^{\Delta N104}$ | $NTD^{\Delta N97}$ | $NTD^{\Delta N54}$ |
| SASBDB Accession Codes | | | | |
| Organism | *Gallus gallus* | | | |
| NCBI protein accession ID | 395427 | | | |
| (amino acid range) | 121-382 | 105-382 | 98-382 | 55-134 |
| SEC-SAXS buffer | 20 mM Tris pH 7.0, 10 mM DTT | | | |
| NaCl concentration | 300 mM | | | |
| Sample injection volume | 40 µl | 40 µl | 40 µl | 40 µl |
| Sample injection conc. | 7.4 mg/ml | 9.5 mg/ml | 6.5 mg/ml | 3.7 mg/ml |
| SEC column | S200 Increase 5/150 | | | |
| SEC flow rate | 0.3 ml/min | | | |
| SEC temperature | 20 °C | | | |
| **Instrument details** | | | | |
| Instrument | EMBL P12 bioSAXS beam line, DESY, Hamburg | | | |
| Exposure time/# frames | 0.25 s (540) | | | |
| X-ray wavelength/energy | 0.124 nm (9996.5 eV) | | | |
| Sample-to-detector distance | 3 m | | | |
| Scattering intensity scale | Arbitrary unit, a.u. | | | |
| SEC-SAXS primary data processing | CHROMIXS (ATSAS 3.0.1) | | | |
| # frames used for averaging | 33 | 28 | 33 | 30 |
| Working s-range ($nm^{-1}$) | 0.15-3.7 | 0.20-2.88 | 0.18-3.17 | 0.08-2.7 |
| **Guinier analysis:** | | | | |
| Primary data analysis software | PRIMUS (ATSAS 3.0.1) | | | |
| Guinier I(0) (σ) | 2046(4) | 5364(7) | 1828(3) | 671(2) |
| $R_g$ (Guinier, nm) (σ) | 2.67(0.01) | 2.68(0.01) | 2.52(0.01) | 2.63(0.01) |
| $sR_g$ range | 0.42-1.15 | 0.67-1.24 | 0.44-1.33 | 0.21-1.30 |
| **p(r) analysis:** | | | | |
| Method | GNOM 5 | | | |
| I(0), POR (σ) | 2074(5) | 5574(9) | 1868(5) | 673(2) |
| $R_g$ (POR, nm) (σ) | 2.86(0-02) | 3.01(0.01) | 2.76(0.02) | 2.68(0.02) |
| $D_{max}$ (nm) | 12.5 | 13.0 | 11.9 | 9.5 |
| Quality of fit, CorMap P / $\chi^2$ | 0.71/1.09 | 0.06/1.12 | 0.29/1.05 | 0.36/1.00 |
| Porod volume ($nm^3$) | 43 | 46 | 42 | 64 |
| Shape classification | extended | extended | compact | flat |
| **Molecular Weight analysis:** | | | | |
| MW, calculated from amino acid sequence, kDa | 29.4 | 31.4 | 32.3 | 37.0 |
| MW from SAXS data, kDa | 37-40 | 25-29 | 31-35 | 29-31 |
| **Ensemble modeling** | | | | |
| Method | EOM | EOM | EOM | EOM |
| Symmetry | P1 | P1 | P1 | P1 |
| Template | PDB ID: 3W9G | PDB ID: 3W9G | PDB ID: 3W9G | PDB ID: 3W9G |
| Final model fit, CorMap P / $\chi^2$ | 1.13e-64/39.45 | 3.01e-66/14.67 | 7.16e-04/1.14 | 0.01/1.02 |

**Table S3 – Lipid composition used in the coarse-grained molecular dynamics simulations to mimic the plasma membrane inner leaflet.**

| Lipid type | MARTINI model | Charge | Abundance [%] |
|---|---|---|---|
| Cholesterol | CHOL | 0 | 20.03 |
| 1-palmitoyl-2-oleoyl-glycero-3-phosphocholine | POPC | 0 | 69.14 |
| 1,2-dioleoyl-glycero-3-phosphoserine | DOPS | -1 | 9.94 |
| 1-palmitoyl-2-linoleoyl-*sn*-glycero-3-phosphoinositol-4,5-bisphosphate | POP2 | -5 | 0.89 |

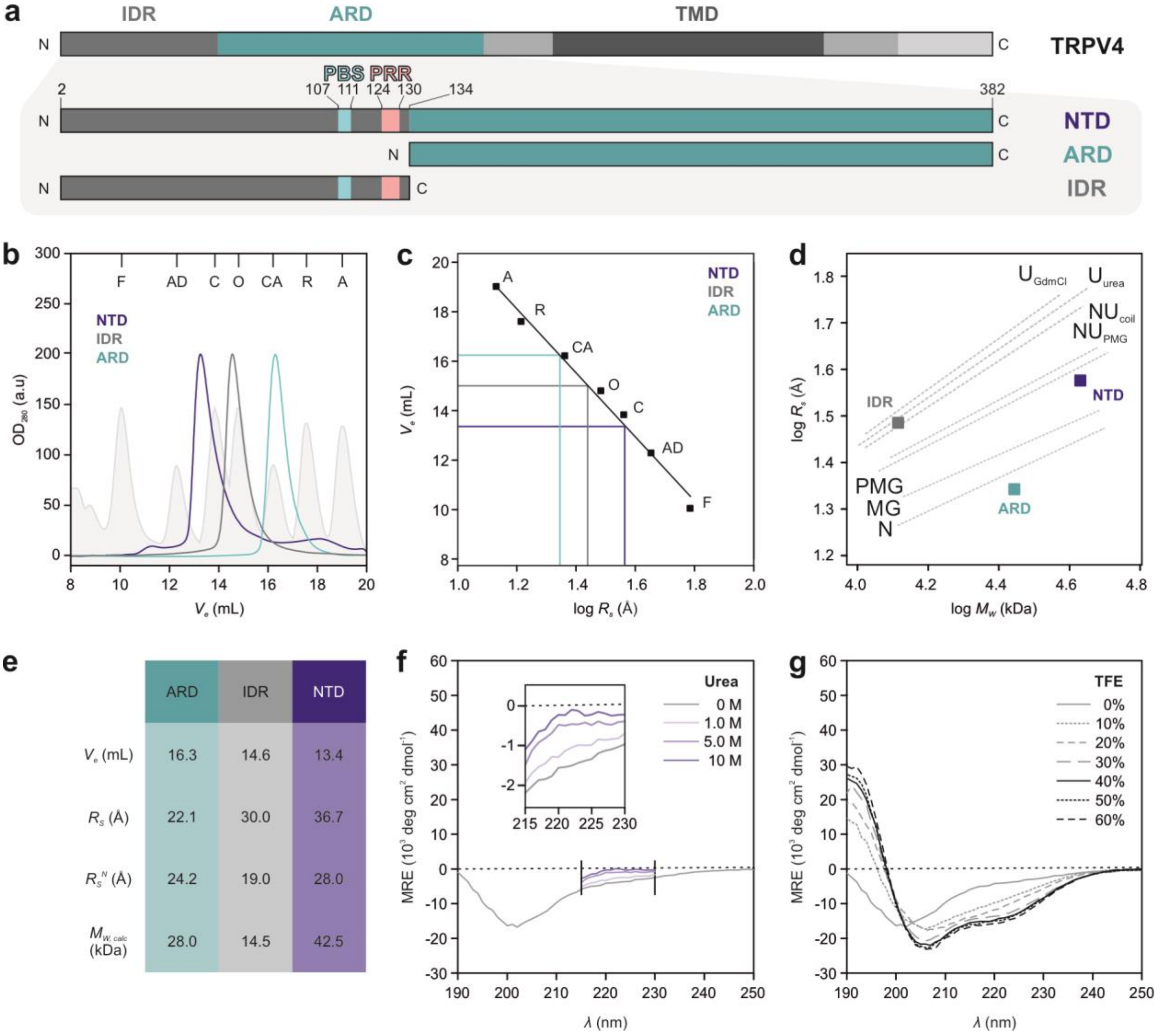


| | ARD | IDR | NTD |
|---|---|---|---|
| $V_e$ (mL) | 16.3 | 14.6 | 13.4 |
| $R_S$ (Å) | 22.1 | 30.0 | 36.7 |
| $R_S^N$ (Å) | 24.2 | 19.0 | 28.0 |
| $M_{W,calc}$ (kDa) | 28.0 | 14.5 | 42.5 |

**Fig. S1: Structural characterization of heterologously expressed TRPV4 N-terminal constructs.**

**a** Construct design for *G. gallus* TRPV4 NTD, ARD and IDR. Important regulatory sites (PBS: $PIP_2$-binding site; PRR: proline rich region) are highlighted in light blue and light pink, respectively.

**b** Analytical size-exclusion chromatography (SEC) of TRPV4 NTD, IDR and ARD. SEC profiles of standard proteins for Stokes radius ($R_S$) determination are shown as filled curves (light grey): F: ferritin; AD: alcohol dehydrogenase; C: conalbumin; O: ovalbumin; CA: carbonic anhydrase; R: ribonuclease A; A: aprotinin. SEC profiles of NTD, IDR and ARD were normalized to an $OD_{280}$ of 200 a.u.

**c** Calibration curve (black) generated with elution volumes of protein standards shown in (b).

**d** Structural analysis of TRPV4 N-terminal constructs based on $M_w$ and newly determined $R_S$. The ARD can be classified as a native, globular protein, the IDR as a native coil-like protein and the NTD as a pre-molten globule-like protein, i.e. a mixture of unfolded and globular states. (N: native; MG: molten globule; PMG: pre-molten globule; $NU_{PMG}$: native pre-molten globule like; $NU_{coil}$: native coil-like; $U_{urea}$: urea unfolded; $U_{GdHCl}$: guanidinium hydrochloride unfolded (for details on classification, see paper by Uversky et al.[1])).

**e** Stokes radii ($R_S$) calculated from the SEC elution volumes ($V_e$) in the calibration curve shown in (d). Theoretical $R_S$ values ($R_S^N$) assuming globular, monomeric proteins under native conditions were determined using described procedures[1] based on the calculated molecular weights ($M_{Wcalc}$) of the three constructs.

**f, g** Far-UV circular dichroism (CD) spectra of purified IDR with increasing amounts of trifluoroethanol (TFE) and urea demonstrates that secondary structure can only be induced at high amounts of TFE and no secondary structure content is lost by urea addition, thereby classifying the IDR as truly intrinsically disordered. Due to high detector voltages in the presence of high urea concentrations, only wavelengths above 215 nm were recorded. The relevant region for changes in secondary structure content is shown in the box.

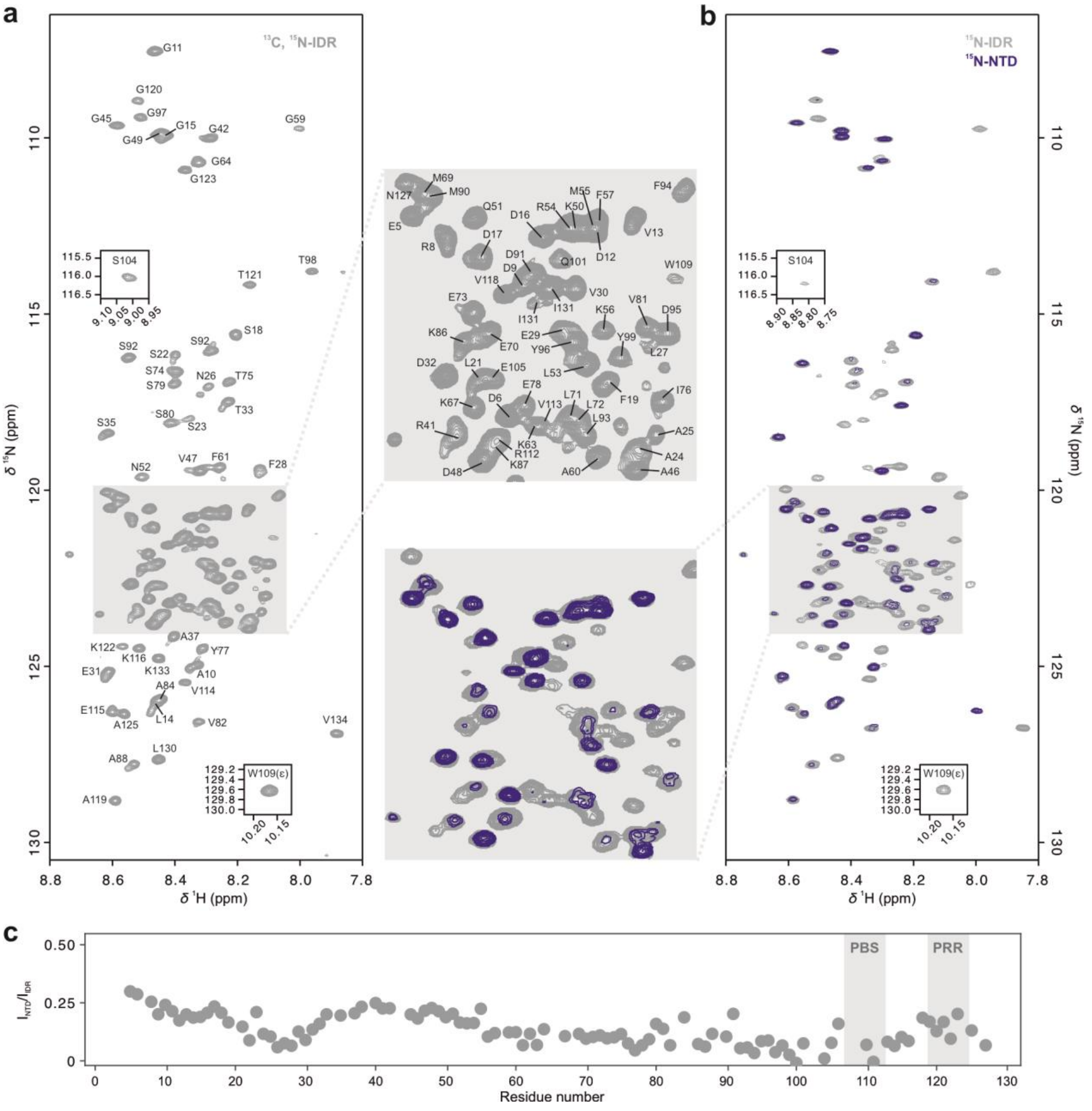

**Fig. S2: [$^1$H, $^{15}$N]-TROSY-HSQC NMR spectra of isolated TRPV4 IDR and in context of the NTD.**
**a** Backbone amide resonance assignments of native *G. gallus* TRPV4 IDR comprising residues 2-134[2].
**b** Overlay of [$^1$H, $^{15}$N]-TROSY-HSQC spectra of $^{15}$N-labeled IDR (grey) and NTD (blue). The resonances of the ARD are not visible, presumably due to unfavorable dynamics (see main text for details).
**c** Relative peak intensity of residues within the IDR in isolation or in the context of the NTD, i.e. in the presence of the ARD. Reduced signal intensity, i.e. a low $I_{NTD}/I_{IDR}$ ratio, suggests intradomain contact sites along the IDR sequence. Previously described functionally important sites in the C-terminal IDR are highlighted by grey boxes (PBS: $PIP_2$-binding site; PRR: proline rich region).

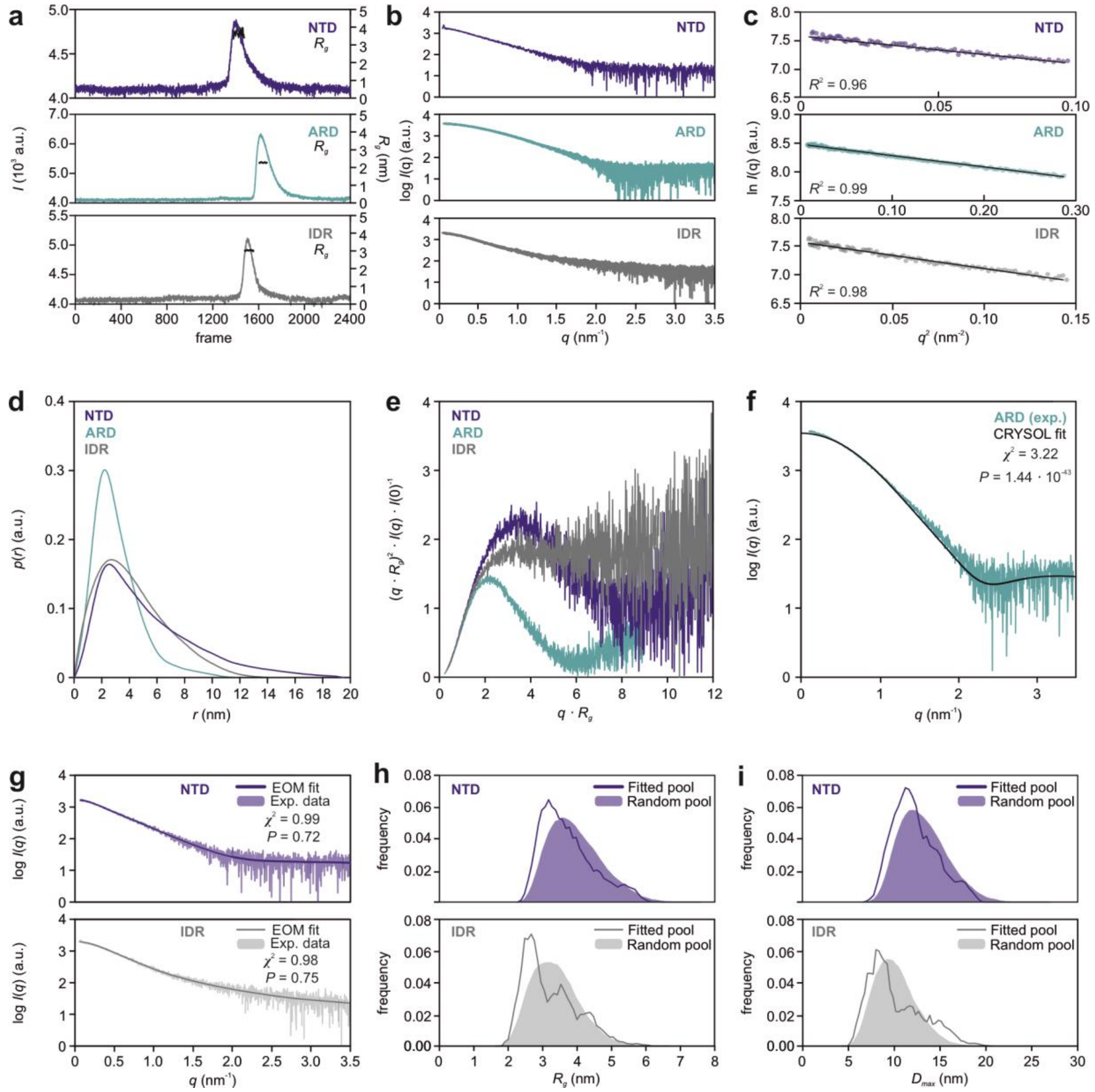


**Fig. S3: SEC-SAXS and Ensemble optimization method (EOM) analysis of TRPV4 N-terminal constructs.**

**a** SEC-SAXS profiles of NTD (blue), ARD (cyan), and IDR (grey). Plotted are the partially integrated scattering intensities, *I*, from sequentially recorded 1D-scattering data frames measured throughout the SEC-elution of each sample. After background/buffer subtraction, the respective radii of gyration ($R_g$) correlation through the elution peaks of each sample were calculated using the Guinier approximation and are indicated in black (with the magnitude on the right axis).

**b** X-ray scattering profiles of N-terminal constructs plotted as the logarithm of the scattering intensity log(*I*(*q*)) (arbitrary units) versus the momentum transfer, *q*.

**c** Guinier-plots (ln*I*(*q*) vs $q^2$, plotted to low-angle: $qR_g < 1.3$) of N-terminal constructs.

**d** Real-space pair-distance distribution functions, or *p*(*r*) profiles calculated N-terminal constructs. p(r) functions were scaled to an area under the curve value of 1.

**e** Dimensionless Kratky plots of N-terminal constructs. The scattering from ARD is consistent with a compact/globular particle, while the IDR is highly flexible. The NTD scattering shares features of both.

**f** CRYSOL fit[3] of isolated ARD shows unsatisfactory match between experimental and predicted scattering curves based on ARD X-ray structure (PDB: 3W9G) as demonstrated by a $\chi^2$ value of 3.22 and a CorMap *P* value of $1.44 \cdot 10^{-43}$.[4]

**g** Fit of EOM refined IDR and NTD volume-fraction weighted model ensembles (solid line) to the experimental SAXS data of NTD (dark blue) and IDR (grey).

**h, i** $R_g$ and $D_{max}$ distributions of the random pool of generated IDR and NTD structures (filled curves) compared to the $R_g$ and $D_{max}$ distributions of the EOM refined ensembles (solid lines).

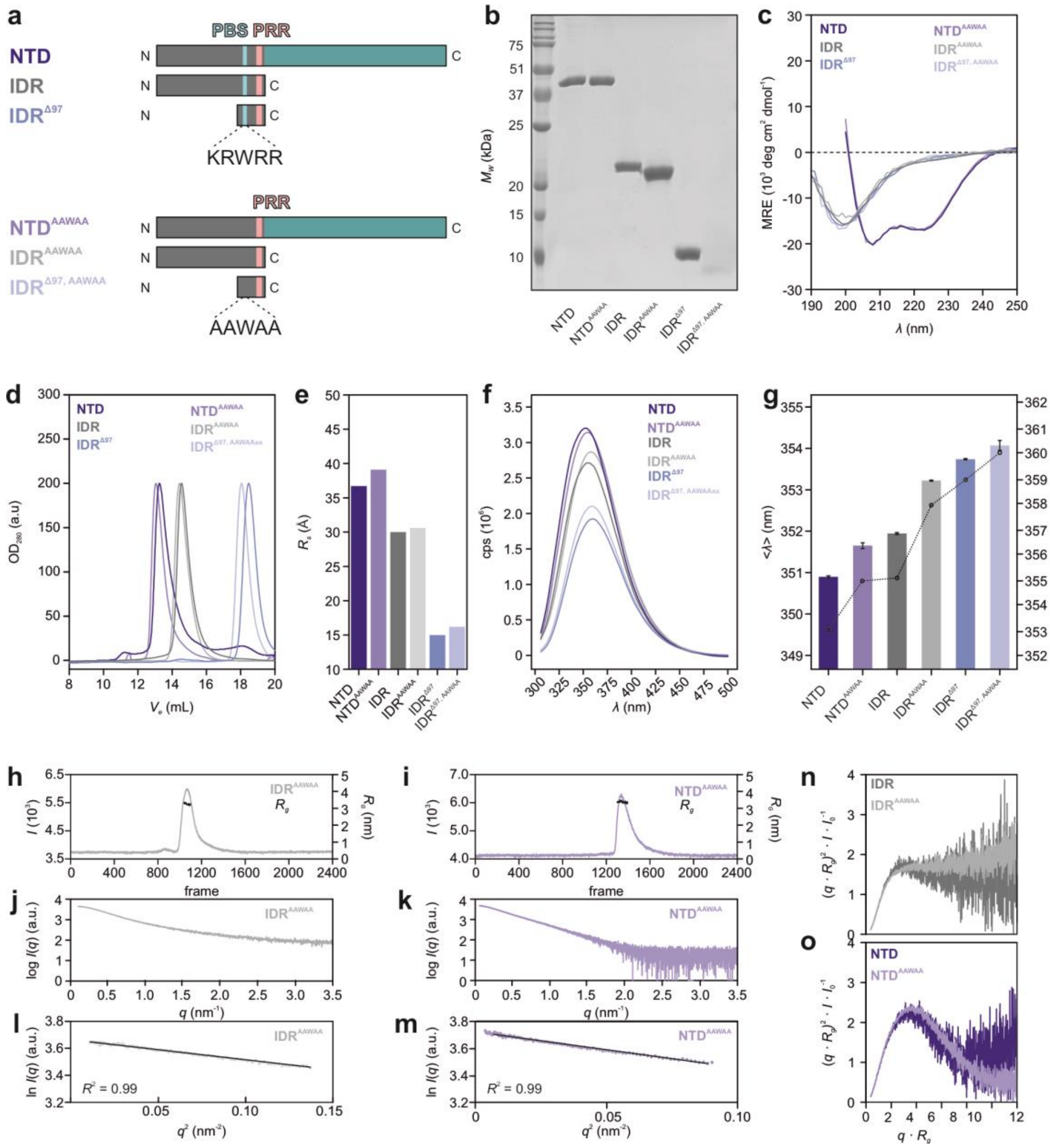


**Fig. S4: PIP$_2$-binding site mutation does not affect the structural integrity of TRPV4 N-terminal constructs.**

**a** Topology of native and PIP$_2$-binding site mutant constructs, NTD, IDR, IDR$^{\Delta N97}$. In the AAWAA mutants, the PIP$_2$-binding site $^{107}$KRWRR$^{111}$ (PBS, yellow) is exchanged to $^{107}$AAWAA$^{111}$ (dark grey). ARD, cyan; IDR, light grey; proline rich region (PRR), light pink.

**b** Coomassie-stained SDS-PAGE of purified native and AAWAA mutant constructs.

**c, d** Far-UV CD spectra (c) and SEC profiles (d) of purified constructs.

**e** Stokes radii ($R_s$) of native and AAWAA mutant constructs estimated from the elution volumes observed in (d).

**f, g** Tryptophan fluorescence spectroscopy of native and corresponding AAWAA mutant constructs. Bars show intensity-weighted average wavelength <$\lambda$> from 320 to 380 nm (left axis). Error bars represent the SD of the mean of n=3. Fluorescence emission maxima $\lambda_{max}$ are shown as black circles connected by dotted lines (right axis).

**h, i** SEC-SAXS profiles of IDR$^{AAWAA}$ (h) and NTD$^{AAWAA}$ (i). Plotted are the partially integrated scattering intensities, *I*, from sequentially recorded 1D-scattering data frames measured throughout the SEC-elution of each sample. After background/buffer subtraction, the respective radii of gyration ($R_g$) correlation through the elution peaks of each sample were calculated using the Guinier approximation and are indicated in black (with the magnitude on the right axis).

**j, k** X-ray scattering profiles of IDR$^{AAWAA}$ (j) and NTD$^{AAWAA}$ (k) plotted as the logarithm of the scattering intensity log($I(q)$) (arbitrary units) versus the momentum transfer, *q*.

**l, m** Guinier-plots (ln$I(q)$ vs $q^2$, plotted to low-angle: $qR_g < 1.3$) of IDR$^{AAWAA}$ (l) and NTD$^{AAWAA}$ (m).

**n , o** Dimensionless Kratky plots of IDR and IDR$^{AAWAA}$ (n) as well as NTD and NTD$^{AAWAA}$ (o).

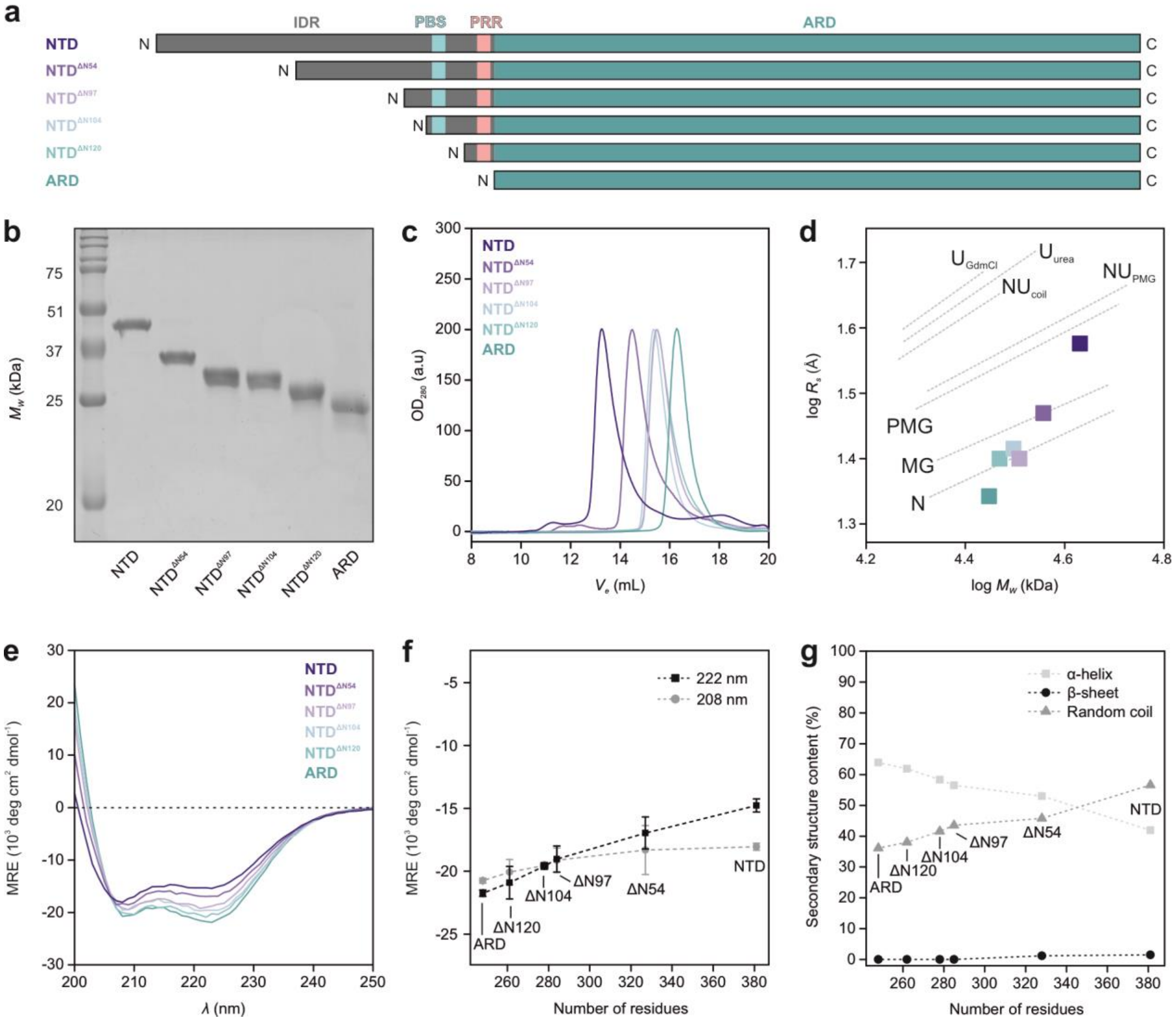


**Fig. S5: Structural characterization of N-terminal TRPV4 NTD deletion mutants.**

**a** N-terminally truncated TRPV4 NTD constructs used in this study.

**b, c** Coomassie-stained SDS-PAGE and SEC profiles of NTD deletion constructs.

**d** $R_s$-$M_W$ analysis of NTD deletion constructs. NTD: $V_e$ = 13.4 mL, $R_s$ = 36.7 Å, $M_W$ = 42.5 kDa; NTD$^{\Delta N54}$: $V_e$ = 14.6 mL, $R_s$ = 29.6 Å, $M_W$ = 37.0 kDa; NTD$^{\Delta N97}$: $V_e$ = 15.4 mL, $R_s$ = 25.4 Å, $M_W$ = 32.3 kDa; NTD$^{\Delta N104}$: $V_e$ = 15.4 mL, $R_s$ = 26.0 Å, $M_W$ = 31.4 kDa; NTD$^{\Delta N120}$: $V_e$ = 15.5 mL, $R_s$ = 25.5 Å, $M_W$ = 29.4 kDa; ARD: $V_e$ = 16.3 mL, $R_s$ = 22.1 Å, $M_W$ = 28.0 kDa.

**e, f** Far-UV CD spectra of NTD deletion constructs and mean residue ellipticity (MRE) at 208 nm and 222 nm. Error bars correspond to SD of mean from n=3.

**g** BeStSel[5] based secondary structure analysis plotted versus the number of amino acids in the NTD constructs shows that overall secondary structure content of the ARD is not affected by consecutive N-terminal deletions of the intrinsically disordered IDR.

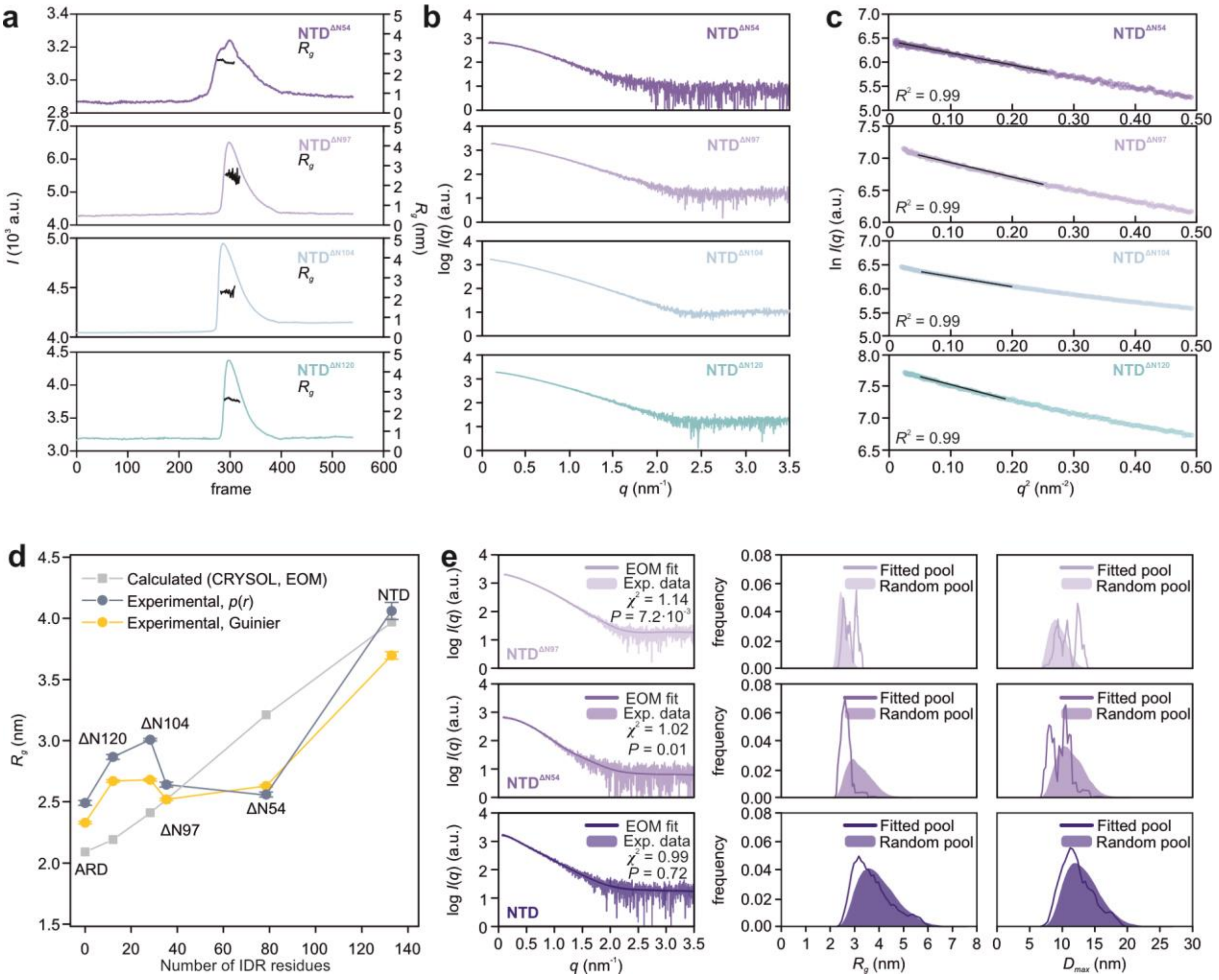


**Fig. S6: SAXS analysis of N-terminal deletion mutants of TRPV4 NTD.**

**a** SEC-SAXS profiles of NTD$^{\Delta N54}$, NTD$^{\Delta N97}$, NTD$^{\Delta N104}$ and NTD$^{\Delta N120}$. Plotted are the partially integrated scattering intensities, *I*, from sequentially recorded 1D-scattering data frames measured throughout the SEC elution of each sample. After background/buffer subtraction, the respective radius of gyration ($R_g$) correlations with the elution peaks of each sample were calculated using the Guinier approximation (indicated in black, right axis).

**b** Background subtracted SAXS profiles of the NTD deletion mutations extracted from the SEC traces shown in (a).

**c** Corresponding Guinier-plot of scattering data shown in (b).

**d** Experimental and calculated radius of gyration ($R_g$) plotted versus the number of IDR residues included in the respective NTD constructs. The $R_g$ of the isolated ARD was calculated with CRYSOL[3] (grey). The $R_g$ of the NTD and the deletion mutants represent the average $R_g$ value of a random pool of conformations calculated via EOM analysis[6] assuming a random chain behavior of the IDR. $R_g$ values were experimentally obtained from the Guinier analysis (yellow) or the pair-distance distribution (blue-grey).

**e** EOM analysis of NTD$^{\Delta N97}$, NTD$^{\Delta N54}$, and wildtype NTD. Shown is the EOM fit (solid line) to the respective experimental data (left). On the right, the EOM derived $R_g$ and $D_{max}$ distributions of a random pool of generated conformations (filled curves) were compared to the EOM-selected ensemble fit to the experimental data. The experimental SAXS profiles of the NTD$^{\Delta N104}$ and NTD$^{\Delta N120}$ mutants could not be reliably fitted to a random pool of conformations using the EOM algorithm (not shown).

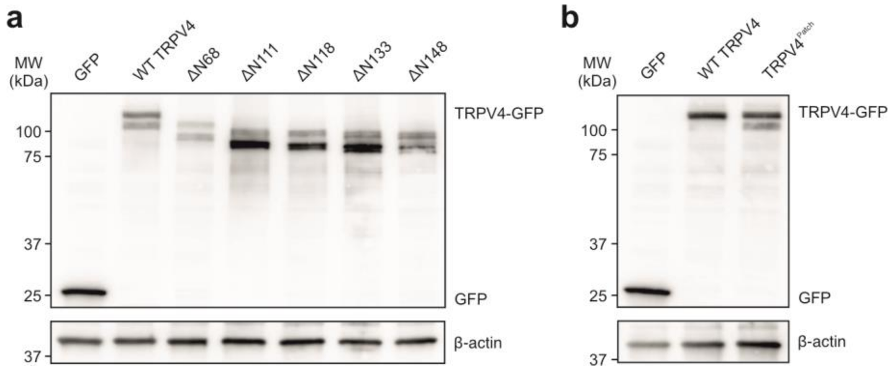


**Fig. S7: Expression of human TRPV4 N-terminal constructs in MN-1 cells.**

**a** Western Blot of consecutive human TRPV4 N-terminal deletion constructs.

**b** Western Blot of human TRPV4$^{Patch}$ mutant.

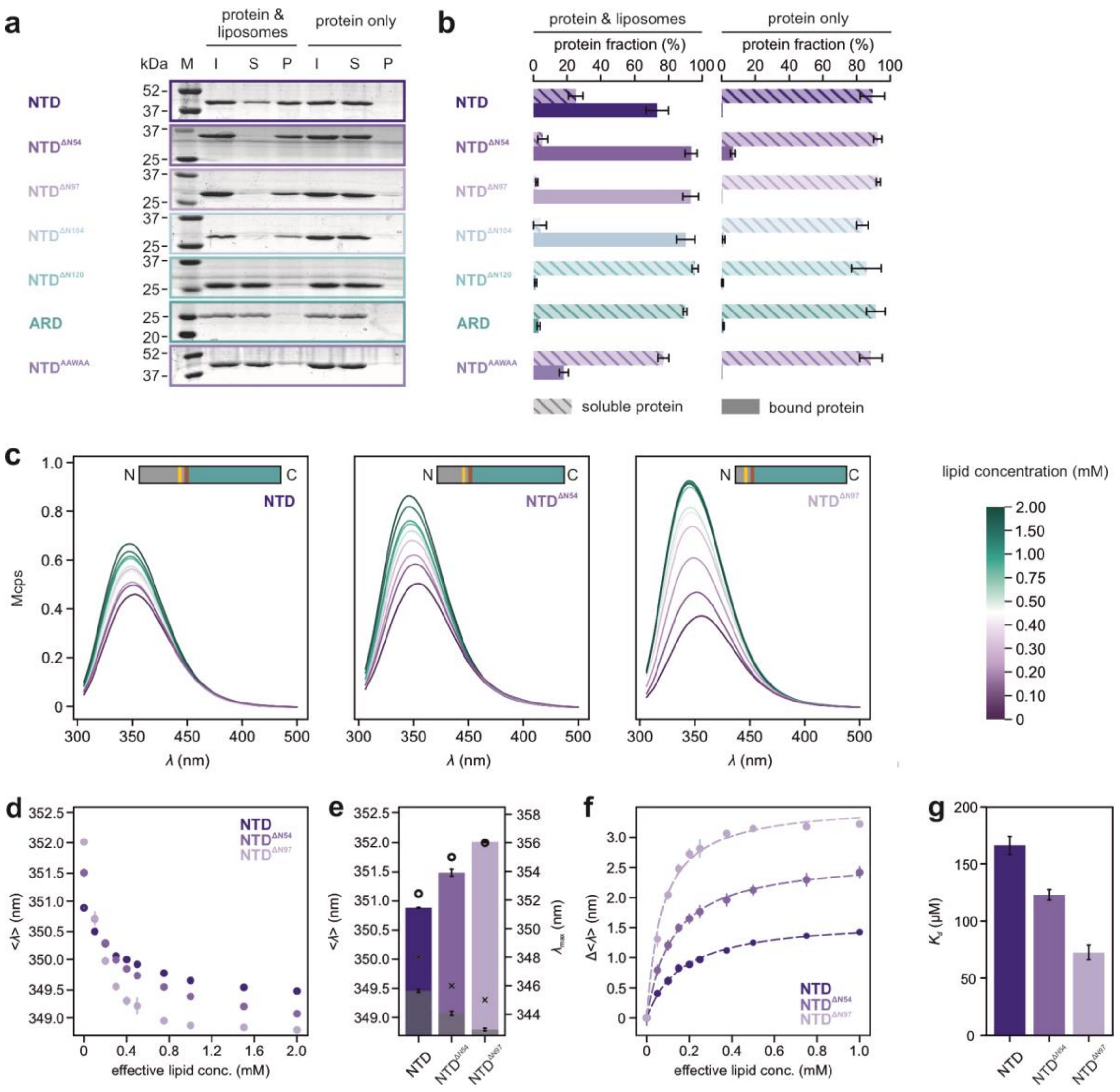


**Fig. S8: Effect of N-terminal truncations on the lipid interaction of the TRPV4 NTD.**

**a** Representative SDS-PAGEs for liposome sedimentation assay. I: input; S: supernatant; P: pellet. Control experiments were carried out in the absence of liposomes ('protein only'). A lipid mixture of 50% (w/w) POPG and 50% (w/w) POPC was used to prepare liposomes with 100 nm diameter.

**b** Protein distribution between pellet and supernatant after centrifugation. The protein fraction in the pellet and supernatant was quantified via densitometry of SDS-PAGE protein bands using imageJ[7]. Representative gels are shown, error bars represent SD of mean from n=3.

**c** Tryptophan fluorescence spectra of TRPV4 NTD and N-terminal deletion mutants in the presence of 0/0.1/0.2/0.3/0.4/0.5/0.75/1.0/1.5/2.0 mM lipids (100 nm liposomes, POPG:POPC in a 1:1 ratio).

**d** Intensity-weighted average wavelength $<\lambda>$ plotted against the lipid concentration.

**e** Intensity-weighted average wavelength $<\lambda>$ of TRPV4 NTD constructs without (colored bars) and in the presence of 2 mM lipid (grey bars) ($<\lambda>$ on left axis). Tryptophan fluorescence emission maximum wavelengths $\lambda_{max}$ without lipids and in the presence of liposomes (2 mM lipid) are shown as black circles and black crosses respectively (values for $\lambda_{max}$ on right axis).

**f** Intensity-weighted average wavelength shift $\Delta<\lambda>$ plotted against the effective lipid concentration (half of total lipid concentration). The data were fitted with a Hill equation with a Hill coefficient of 1 to obtain $K_d$ values.

**g** Comparison of determined $K_d$ values in (d) for NTD (165.1 ± 7.9 µM), NTD$^{\Delta N54}$ (122.1 ± 4.6 µM) and NTD$^{\Delta N97}$ (72.1 ± 6.4 µM).

The error bars in (b, c, d) represent the SD of mean from n=3. The error bars in (e) are the fit errors from (d).

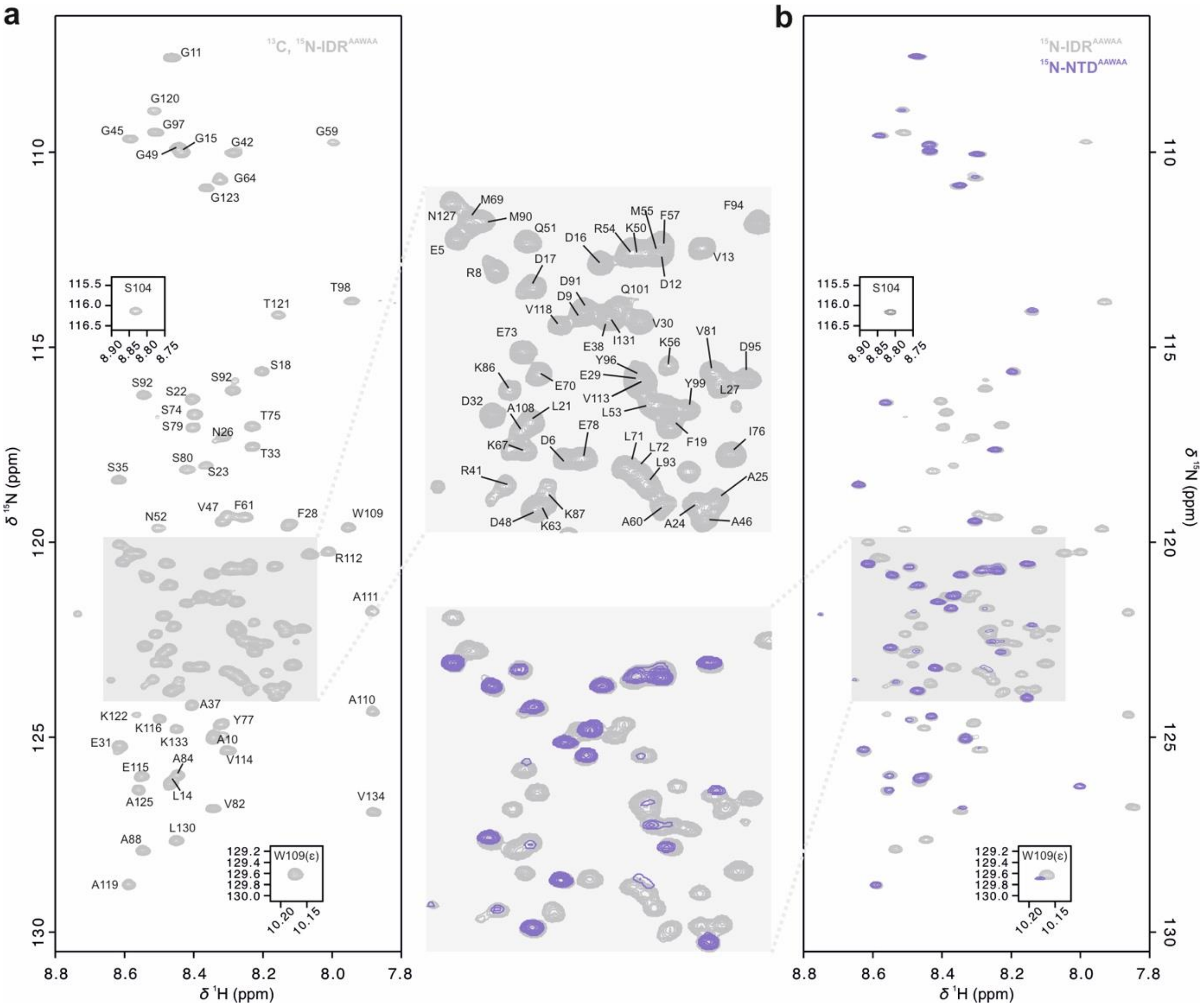


**Fig. S9: NMR backbone assignments of TRPV4 IDR$^{AAWAA}$.**

**a** [$^{1}$H, $^{15}$N]-TROSY-HSQC NMR spectrum of $^{13}$C, $^{15}$N-labeled IDR$^{AAWAA}$. The basic residues in the PIP$_2$-binding site ($^{107}$KRWRR$^{111}$) have been mutated to alanine.

**b** Overlay of the spectra of $^{15}$N-labeled IDR$^{AAWAA}$ and NTD$^{AAWAA}$.

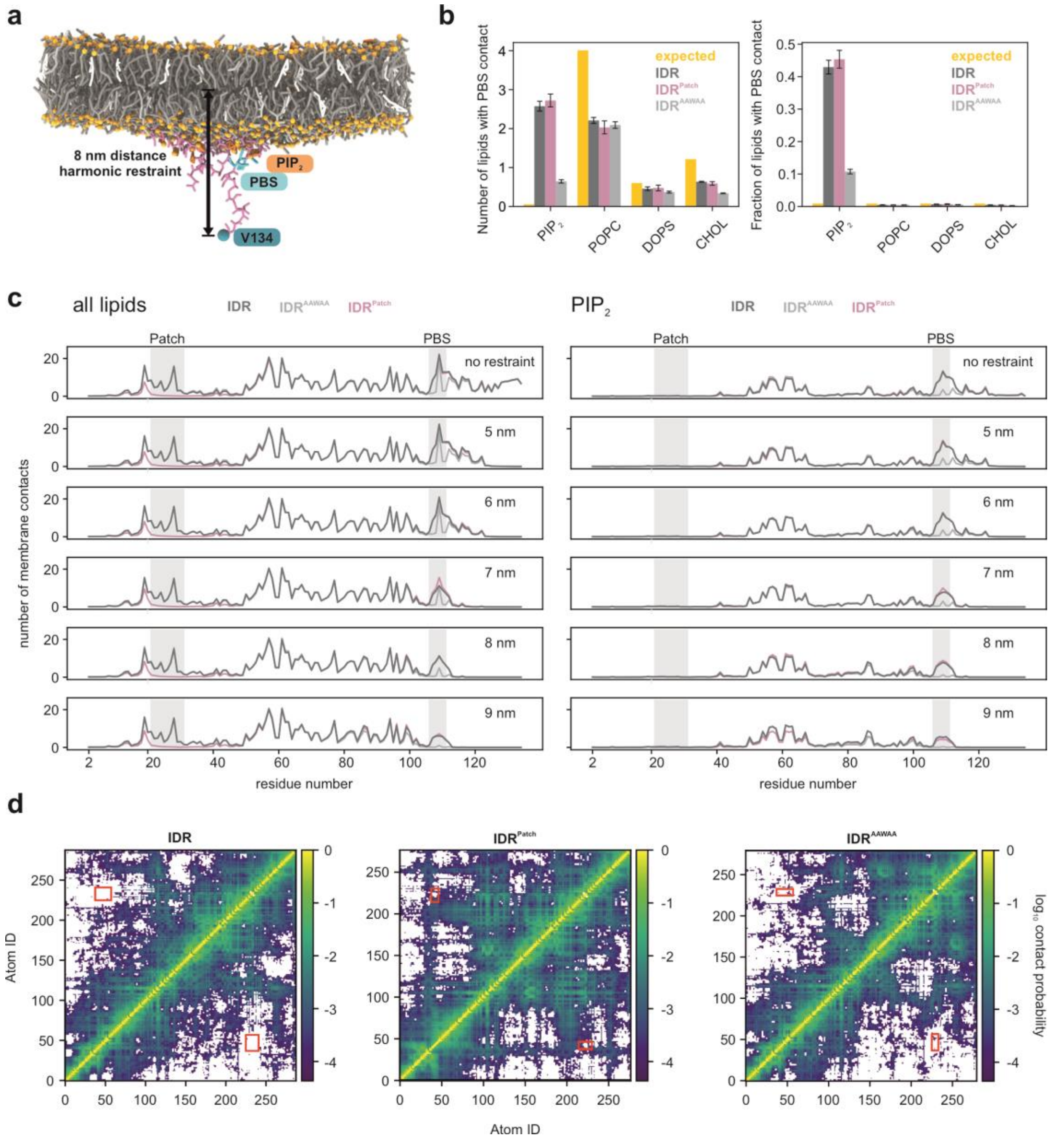


**Figure S10: TRPV4 IDR interactions with the plasma membrane in coarse-grained MD simulations.**

**a** Snapshot of the bent membrane from a simulation of the native IDR (pink) restrained at 8 nm (distance to membrane midplane) after 15.5 µs of simulation on a lipid bilayer membrane consisting of $PIP_2$ (1%, dark orange), POPC (69%, dark grey), DOPS (10%, light gray) and cholesterol (20%, white) (see Table S3). Headgroup phosphates are shown as orange spheres. The $PIP_2$-binding site of the TRPV4 IDR is highlighted in cyan, the C-terminal residue V134 as a blue sphere.

**b** Absolute (left) and relative (right) number of contacts between the $PIP_2$-binding site (PBS) and each lipid type in simulations of the native IDR (dark grey), $IDR^{AAWAA}$ (light grey) and $IDR^{Patch}$ (mauve). Values are based on simulations carried out without a height restraint. Yellow bars indicate the number of expected contacts for the $PIP_2$-binding site based on the respective mol-fractions of lipid types and the observed total number of contacts. Error bars depict the standard error of the mean (SEM) of the individual replicate simulations.

**c** Average number of membrane contacts for each IDR residue for all lipids (left) or $PIP_2$ (right). The distance between residue V134 in the IDR C-terminus and the membrane midplane was either not restrained (top panels) or restrained at a specific height (5-9 nm, bottom panels) to emulate the role of the ARD on IDR positioning.

For all conditions shown in (a-c), four replicate simulations per IDR sequence were carried out for ~38 µs and contact averages were calculated from the last ~28 µs of each simulation.

**d** Heat map showing the probability of two coarse-grained atoms of the IDR being in contact with each other in simulations of the native IDR (left), the $IDR^{Patch}$ (center) and the $IDR^{AAWAA}$ (right). Contact maps were calculated after pooling the last ~28 µs of all four replicate simulations without height restraints. A contact was counted when two coarse-grained atoms came closer than 0.6 nm. Red boxes indicate the area of N-terminal patch interaction with the PBS.

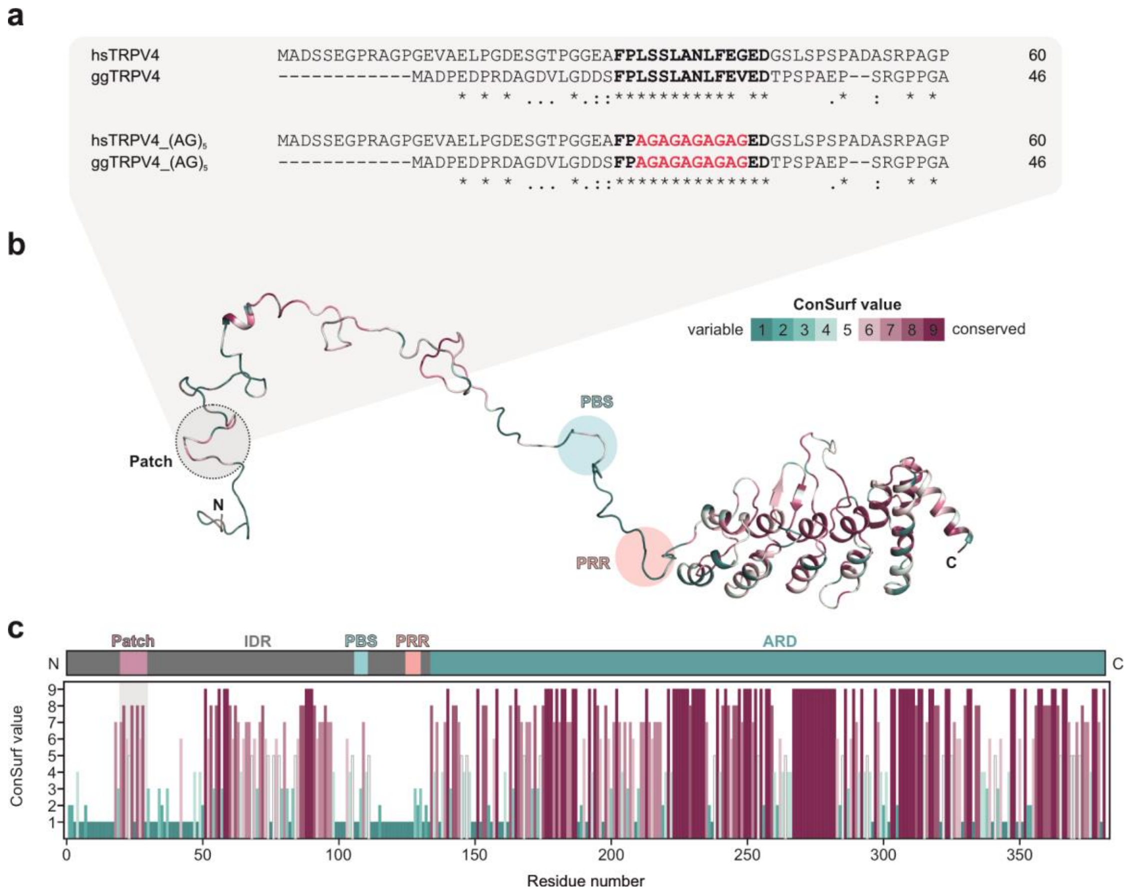


**Fig. S11: sequence alignment and conservation of *H. sapiens* and *G. gallus* TRPV4 IDR.**

**a** Sequence alignment of N-terminal IDR of human and chicken TRPV4. The conserved regulatory patch is shown in bold. The corresponding $(AG)_5$ mutants of the conserved patch are shown below and highlighted in red. Alignments were carried out with ClustalOmega[8].

**b** Conservation of residues in TRPV4 NTD shown mapped onto a *G. gallus* TRPV4 NTD conformer obtained from our EOM analysis (Fig. 1). Analysis was carried out with the ConSurf server[9] (PBS: $PIP_2$-binding site; PRR: proline rich region).

**c** TRPV4 NTD topology and sequence conservation of TRPV4 NTD residues based on the ConSurf analysis[9].

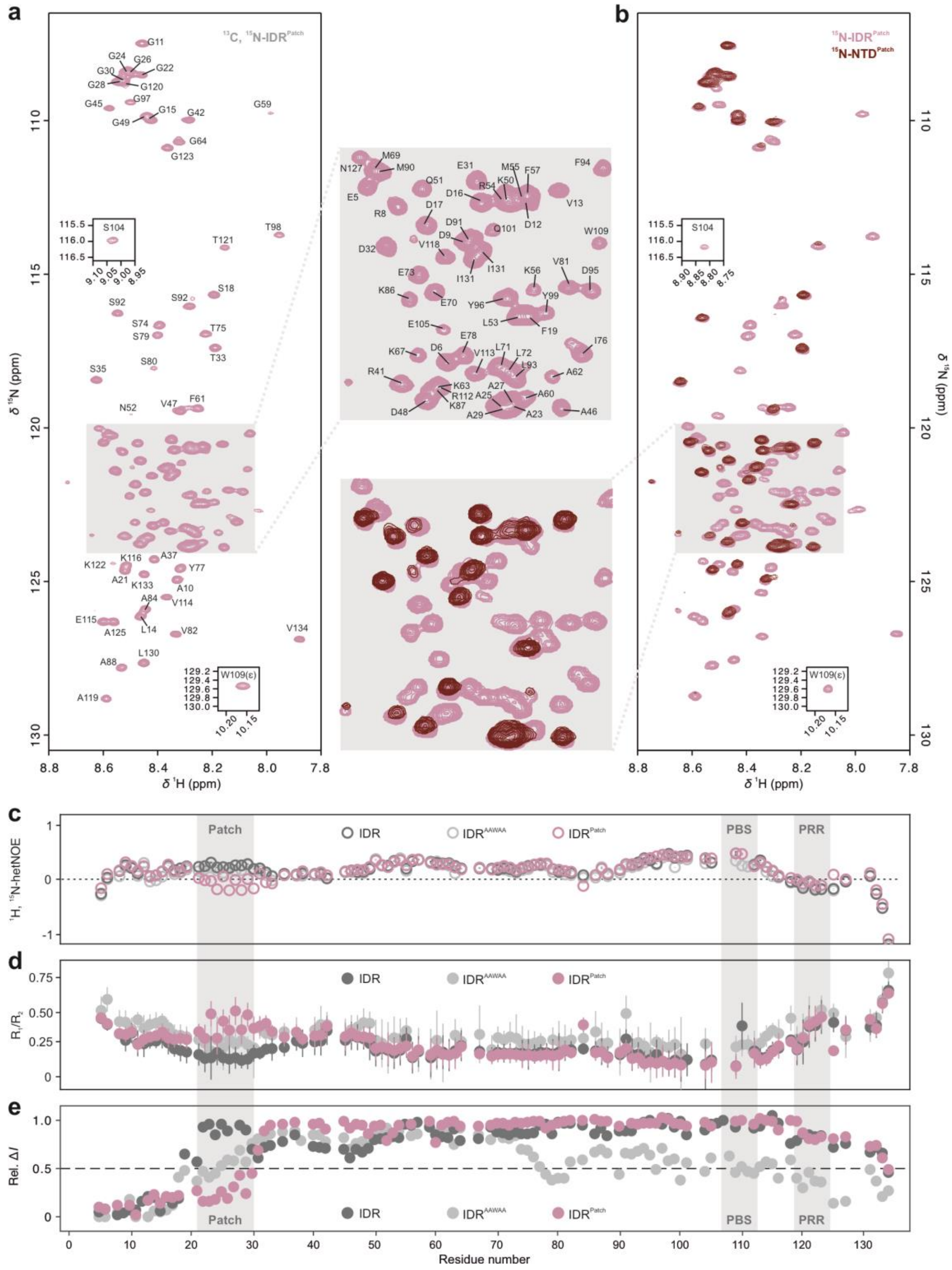


**Fig. S12: Backbone NMR assignment of TRPV4 IDR$^{Patch}$, relaxation measurements of IDR mutants and liposome titrations.**

**a** [$^{1}$H, $^{15}$N]-TROSY-HSQC NMR spectrum of $^{13}$C, $^{15}$N-labeled IDR$^{Patch}$. Here, residues $^{19}$FPLSSLANLFEVE$^{31}$ were exchanged to glycine and alanine yielding $^{19}$FP(AG)$_5$E$^{31}$ (see Fig. S11).

**b** Overlay of the HSQC spectra of $^{15}$N-labeled IDR$^{Patch}$ and NTD$^{Patch}$.

**c, d** {$^{1}$H}$^{15}$N-hetNOE and T$_1$/T$_2$ relaxation data of of $^{15}$N-labeled IDR (dark grey), IDR$^{AAWAA}$ (light grey), IDR$^{Patch}$ (mauve).

**e** NMR signal intensity differences for $^{15}$N-labeled IDR (dark grey), IDR$^{AAWAA}$ (light grey) and IDR$^{Patch}$ (mauve) in the absence and presence of POPC-POPG containing liposomes. For better comparison, the data from Fig. 6d, e for native IDR and IDR$^{AAWAA}$ were added here.

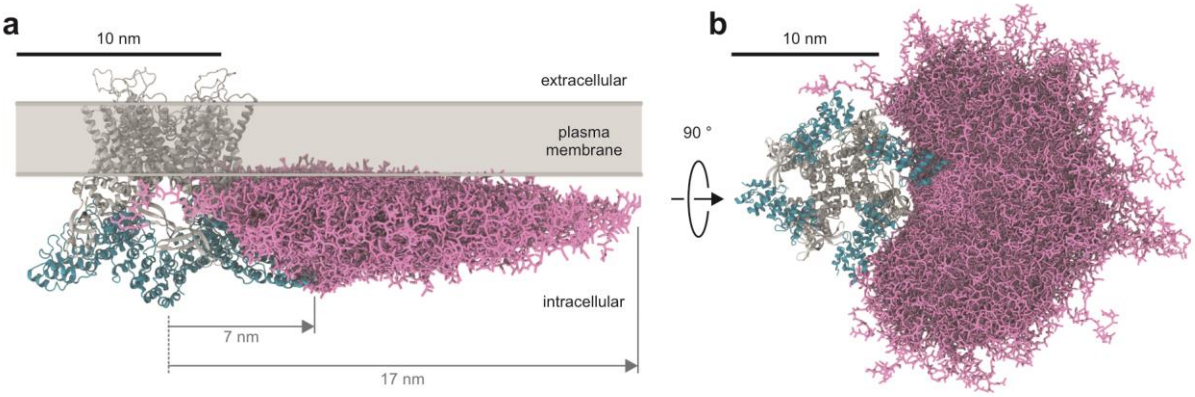


**Figure S13: Structural model of the conformational space sampled by the IDR of a single TRPV4 subunit in the presence of the plasma membrane.**
**a, b** Superimposed IDR conformations (pink licorice) from coarse-grained MD simulations shown for one subunit in the TRPV4 tetramer from the side (a) and top (b). The structured transmembrane core (grey) including the ARDs (cyan) of *G. gallus* TRPV4 has been predicted by AlphaFold multimer[10].

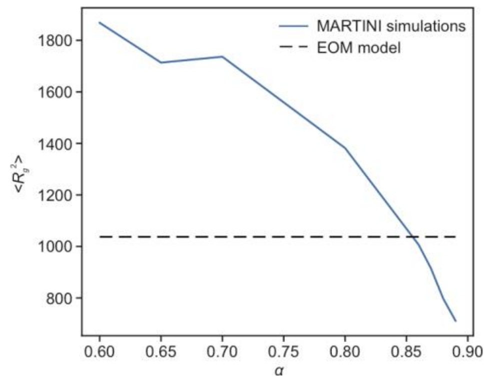


**Fig. S14: Optimizing the scaling factor in MD simulations to best describe the measured $R_g$ distribution of the native IDR.**

Averaged squared $R_g$ ($<R_g^2>$) in simulations with different rescaling constants of the protein/protein interactions ($\alpha$). All simulations were performed without a membrane. The dotted black line shows the averaged squared $R_g$ of the EOM refined model (Fig. 1, Fig. S. 3)

#### Supporting Information References
